# A Model-Based Clustering via Mixture of Bayesian Hierarchical Models with Covariate Adjustment for Detecting Differentially Expressed Genes from Paired Design

**DOI:** 10.1101/2022.02.16.480754

**Authors:** Yixin Zhang, Wei Liu, Weiliang Qiu

## Abstract

The causes of many complex human diseases are still largely unknown. Genetics plays an important role in uncovering the molecular mechanisms of complex human diseases. A key step to characterize the genetics of a complex human disease is to unbiasedly identify disease-associated gene transcripts in whole-genome scale. Confounding factors could cause false positives. Paired design, such as measuring gene expression before and after treatment for the same subject, can reduce the effect of known confounding factors. Model-based clustering, such as mixtures of Bayesian hierarchical models, has been proposed to detect gene transcripts differentially expressed between paired samples. However, not all known confounding factors can be controlled in a paired/match design. To the best of our knowledge, no clustering methods have the capacity to adjust for the effects of covariates yet. In this article, we proposed a novel mixture of Bayesian hierarchical models with covariate adjustment in identifying differentially expressed transcripts using high-throughput whole-genome data from paired design. Both simulation study and real data analysis show the good performance of the proposed method

## Introduction

Genome-wide differential gene expression analysis is widely used for the elucidation of the molecular mechanisms of complex human diseases. One popular and powerful approach to detect differentially expressed genes is the probe-wise linear regression analysis combined with the control of multiple testing, such as limma^1^. That is, we first perform linear regression for each probe and then adjust p-values for controlling multiple testing. One advantage of this approach is its capacity to adjust for potential confounding factors.

Another approach for detecting differentially expressed genes is the model-based clustering via mixture of Bayesian hierarchical model (MBHM)^2–7^, which can borrow information across genes to cluster genes. Probe clustering based on MBHMs treats gene transcripts as “samples” and samples as “variables”. Therefore, transcript clustering based on MBHMs has large number of “samples” and relatively small number of “variables”, hence does not have the curse-of dimensionality problem. In addition, unlike transcript-specific tests that have several parameters per transcript, transcript clustering based on MBHMs has only a few hyperparameters per cluster to be estimated and could borrow information across transcripts to estimate model hyperparameters. These approaches generally assume that samples under two groups are obtained independently.^8^ proposed a constrained MBHM to identify genetic outcomes measured from paired/matched designs.

Paired design is commonly used in study design for its homogeneous external environment for comparing measurements under different conditions. However, not all known confounding factors can be controlled in a paired/match design. Hence, we might still need to adjust the effects of confounding factors for data from a paired/matched design. To best of our knowledge, no existing clustering methods, including the constrained MBHM^8^, have the capacity to adjust for potential confounding factors, i.e., age, sex, and/or batch effect.

In this article, we proposed a novel mixture of Bayesian hierarchical models with covariate adjustment in identifying differentially expressed transcripts using high-throughput whole genome data from paired design.

## Materials and Methods

We assumed that gene transcripts can be roughly classified into 3 clusters based on their expression levels in subjects after treatment (denoted as condition 1) relative to those before treatment (denoted as condition 2):

1. transcripts after treatment have higher expression levels than those before treatment, i.e., over-expressed (**OE**) in condition 1;
2. transcripts after treatment have lower expression levels than those before treatment, i.e., under-expressed (**UE**) in condition 1;
3. transcripts after treatment have same expression levels than those before treatment, i.e., non-differentially expressed (**NE**) between condition 1 and matched condition 2.

We followed^8^ to directly model the marginal distributions of gene transcripts in the 3 clusters. In^8^, they proposed a mixture of three-component Bayesian hierarchical distributions to characterize the within-pair difference of gene expression. We extended their model by incorporating potential confounding factors (such as *Age* and *Sex*) in the mixture of Bayesian hierarchical models, which might affect the response of gene expression to drug treatment. We assumed that data have been processed so that the distributions of mRNA expression levels are close to normal distributions. For RNAseq data, we can apply VOOM transformation^9^ or countTransformers^10^.

### A mixture of Bayesian hierarchical models

For the *g*^*th*^ gene transcript, let *x*_*gl*_ and *y*_*gl*_ denote the expression levels of the *l*^*th*^ subject under two different conditions, e.g., before and after treatment, *g* = 1, …, *G, l* = 1, …, *n*, where *G* is the number of transcripts and *n* is the number of subjects (i.e., the number of pairs). Let *d*_*gl*_ = log_2_(*y*_*gl*_) − log_2_(*x*_*gl*_) be the log2 differences for the *l*^*th*^ subject. Denote *d*_*g*_ = (*d*_*g*1_, …, *d*_*gn*_)^*T*^. We assumed that *d*_*gl*_ is conditionally normally distributed given mean vector and covariance matrix. Let *X*^*T*^ be the *n*× (*p* + 1) design matrix, where *p* is the number of covariates. The first column of *X*^*T*^ is the vector of ones, indicating intercept. Let *η* be the (*p* + 1) ×1 vector of coefficients for the intercept and covariate effects. We assume following mixture of three-component Bayesian hierarchical models:

For the OE gene transcripts,

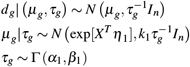

where *k*_1_ > 0, *α*_1_ > 0 and *β*_1_ > 0. Γ(*α*_1_, *β*_1_) denotes the Gamma distribution with shape parameter *α*_1_ and rate parameter *β*_1_. That is, we assume that (1) the mean vectors *μ*_*g*_, *g* = 1, …, *G*, given the variance 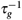 follow a multivariate normal distribution with mean vector exp [*X*^*T*^ *η*_1_] and covariance matrix 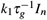; and (2) the variances 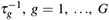, follow a Gamma distribution with shap parameter *α*_1_ and rate parameter *β*_1_.

For the UE gene transcripts,

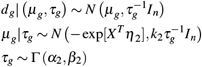

where *k*_2_ > 0, *α*_2_ > 0, *β*_2_ > 0, and *X*^*T*^ is the design matrix.

For the NE gene transcripts,

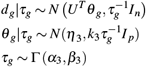

where *k*_3_ > 0, *α*_3_ > 0 and *β*_3_ > 0. *U*^*T*^ is the design matrix without intercept column.

The hyper-parameters *α*_*c*_ and *β*_*c*_ are shape and rate parameters for the Gamma distribution, respectively, *c* = 1, 2, 3. As for *k*_1_, *k*_2_ and *k*_3_, the variation of the mean vector *μ*_*g*_ should be smaller than that of the observations *d*_*g*_. So we expect 0 *< k*_*c*_ *<* 1, *c* = 1, 2, 3.

Note that the marginal distribution for each component of the mixture is a multivariate t distribution with special structure of mean vector and covariance matrix [11, Section 3.7.6].

For continuous covariates, we require that they are standardized so that they have mean zero and variance one. Standardizing continuous covariates would make exp (*X*^*T*^ *η*_1_) and exp (*X*^*T*^ *η*_2_) be numerically finite.

Ideally, we should require *μ*_*g*_ > 0(*μ*_*g*_ *<* 0) for all transcripts in cluster 1 (cluster 2). To do so, we can assume a log normal prior distribution for *μ*_*g*_ in cluster 1, for instance. However, a log normal distribution could not be a conjugate prior for the mean of a normal distribution. It would increase the computational burden if non-conjugate priors were used. As an alternative, we require the mean *E*(*μ*_*g*_) > 0 (*E*(*μ*_*g*_) *<* 0) for cluster 1 (cluster 2) by assuming *E*(*μ*_*g*_) for cluster 1 (cluster 2) to be exp[*X*^*T*^ *η*_1_] (−exp[*X*^*T*^ *η*_2_]). Denote 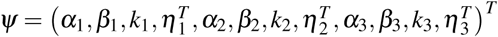.

### Marginal density functions

Let *f*_1_(**d**_*g*_ |*ψ*), *f*_2_(**d**_*g*_ |*ψ*), *f*_3_(**d**_*g*|_ *ψ*) be the marginal densities of the 3 Bayesian hierarchical models, and *π* = (*π*_1_, *π*_2_, *π*_3_) is the vector of cluster mixture proportions. Then the marginal density of **d**_*g*_ is :

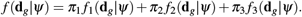

### Determining transcript cluster membership

The transcript-cluster membership is determined based on the posterior probabilities, *ζ*_*gc*_ = *Pr*(*g*^*th*^ gene transcript in cluster *c* |*d*_*g*_). We can get

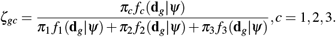

We determine a transcript’s cluster membership as follows: If the maximum value among *ζ*_*gi*_, *i* = 1, 2, 3 is *ζ*_*gc*_, then the transcript *g* belongs to cluster *c*.

The true values of *π*_1_, *π*_2_, *π*_3_, and *ψ* are unknown. We use estimated values to determine transcripts’ cluster membership.

### Parameter estimation via EM algorithm

We used expectation–maximization (EM) algorithm^12^ to estimate the model parameters *π* = (*π*_1_, *π*_2_, *π*_3_)^*T*^ and *ψ*. Detailed derivations of the expectation *Q* of the log complete-data likelihood function and its first-order derivatives are shown in the Appendix.

We stop the expectation-maximization iterations if the sum of squared differences of model parameter estimates between current iteration and previous iteration is smaller than a small constant (e.g. 1.0 ×10^−3^).

More details about the EM algorithm are shown in Appendix.

### A real data study

We downloaded the dataset GSE24742^13^ from the Gene Expression Omnibus (https://www.ncbi.nlm.nih.gov/geo/query/acc.cgi?acc=GSE24742) to evaluate the performance of the proposed model-based clustering method (denoted as *eLNNpairedCov*).

The dataset is from a study that investigated the gene expression before and after administrating rituximab, a drug for treating anti-TNF resistant rheumatoid arthritis (RA). There are 12 subjects, each having 2 samples (one sample is before treatment and the other is after treatment). Age and sex are also available. Expression levels of 54,675 gene probes were measured for each of the 24 samples by using Affymetrix HUman Genome U133 Plus 2.0 array. The dataset has been preprocessed by the dataset contributor. We further kept only 43,505 gene probes in the autosomal chromosomes (i.e., chromosomes 1 to 22). We then performed log2 transformation for gene expression levels. We next obtained the within-subject difference of the log2 transformed expression levels (log2 expression after-treatment minus log2 expression before-treatment). By examining the histogram (Figure 5) of the estimated standard deviations of log2 differences of within-subject gene expression for the 43,505 gene probes, we found a bimodal distribution. We excluded gene probes having standard deviation *<* 1. Finally, 23,948 gene probes kept in the down-stream analysis.

### A simulation study

We performed a simulation study to compare the performance of the proposed method *eLNNpairedCov* with transcript-wise test *limma* and Li et. al.’s^8^ method (denoted as *eLNNpaired*). Both *eLNNpairedCov* and *limma* adjust covariate effects, while *eLNNpaired* does not.

The *limma* approach first performs an empirical-Bayes-based linear regression for each transcript. In this linear regression, the within-subject log2 difference of transcript expression is the outcome and intercept indicating if the transcript is over- expressed (intercept>0), under-expressed (intercept*<*0), or non-differentially expressed (intercept = 0), adjusting for potential confounding factors. Here condition 1 are observations after treatment and condition 2 are observations before treatment. A transcript is claimed as OE if its intercept estimate is positive and corresponding FDR-adjusted p-value *<* 0.05, where FDR stands for false discovery rate. A transcript is claimed as UE if its intercept estimate is negative and corresponding FDR-adjusted p-value *<* 0.05. Other transcripts are claimed as NE.

In this simulation study, we considered two scenarios. In the first scenario (Scenario1), the number of subjects is equal to 30. In the second scenario (Scenario2), the number of subjects is equal to 100.

For each scenario, we generated 100 datasets. Each simulated dataset contains *G* = 20, 000 gene transcripts, 0.0376% of which are OE and 0.346% of which are UE.

There are two covariates: standardized age (denoted as *Age*.*s*) and *Sex. Age*.*s* follows normal distribution with mean 0 and standard deviation 1. Seventy five percent (75%) of subjects are women.

The model hyperparameters are set as *α*_1_ = 3.53, *β*_1_ = 3.45, *k*_1_ = 0.26, *η*_1_ = (0.18, 0.00, − 1.05)^*T*^, *α*_2_ = 3.53, *β*_2_ = 3.45, *k*_2_ = 0.26, *η*_2_ = (0.18, 0.00, −1.05)^*T*^, *α*_3_ = 2.86, *β*_3_ = 2.20, *k*_3_ = 0.72, and *η*_3_ = (− 0.01, 0.00)^*T*^.

The parameter values (*π, ψ*, and proportion of women) in the simulation study are estimated from the analysis of the pre-processed real dataset GSE24742 described in Section.

### Evaluation criteria

Two agreement indices and two error rates are used to compare the predicted cluster membership and true cluster membership of all genes. The two agreement indices are accuracy (i.e., proportion of predicted cluster membership equal to the true cluster membership) and Jaccard index^14^. For perfect agreement, these indices have a value of one. If an index takes a value close to zero, then the agreement between the true transcript cluster membership and the estimated transcript cluster membership is likely due to chance. The two error rates are false positive rate (FPR) and false negative rate (FNR). FPR is the percentage of detected DE transcripts among truly NE transcripts. FNR is the percentage of detected NE transcripts among truly DE transcripts. We also examined the user time and number of EM iterations for running each simulated dataset.

## Results

### Results of the real data analysis

For the real dataset, we adjusted standardized age and sex for *eLNNpairedCov* and *limma*. We standardized age so that it has mean zero and variance one. For each transcript, we also scaled its expression across subjects so that its variance is equal one.

The *limma* method detected 6 under-expressed gene transcripts (Figure 1 and Table 1), while *eLNNpaired* did not find any positive signals (i.e., 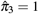). The proposed method *eLNNpairedCov* detected 9 OE transcripts (Upper panel of Figure 2 and Table 2) and 83 UE transcripts (Lower panel of Figure 2 and Table 3). The 83 UE transcripts detected by *eLNNpairedCov* include the 6 UE transcripts detected by *limma*.

**Figure 1.**
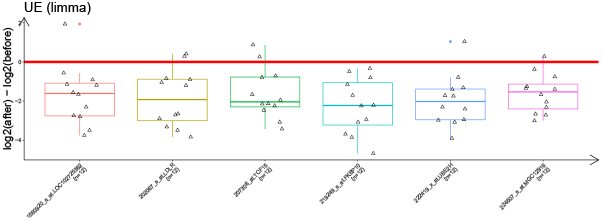
Parallel boxplots of log2 within-subject difference of gene expression for 6 UE transcripts detected by *limma* for pre-processed GSE24742 dataset. Red horizontal line indicates log2 difference equal to zero.

**Figure 2.**
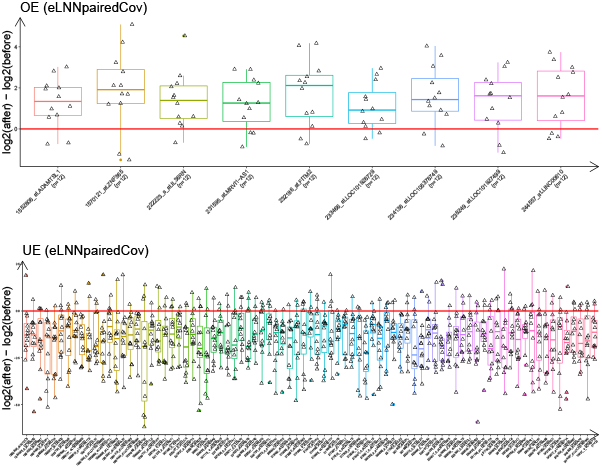
Parallel boxplots of log2 within-subject difference of gene expression for differentially expressed transcripts detected by *eLNNpairedCov* for pre-processed GSE24742 dataset. Upper level: 9 OE transcripts; Lower panel: 83 UE transcripts. Red horizontal lines indicate log2 difference equal to zero.

It is assuring that several genes corresponding to the 92 DE transcripts identified by *eLNNpairedCov* have been associated to rheumatoid arthritis (RA) in literature. For example, Humby et. al. (2019)^15^ reported that genes *ZNF365* (OE), *IL36RN* (OE), *MRVI1-AS1* (OE), *WFDC6* (UE), *UBE2H* (UE), are associated with RA.

### Results of the simulation study

The results for Scenario 1 (*n* = 30) are shown in Figures 3 and 4, which are jittered scatter plots of the performance indices and the difference of performance indices versus methods, respectively.

**Figure 3.**
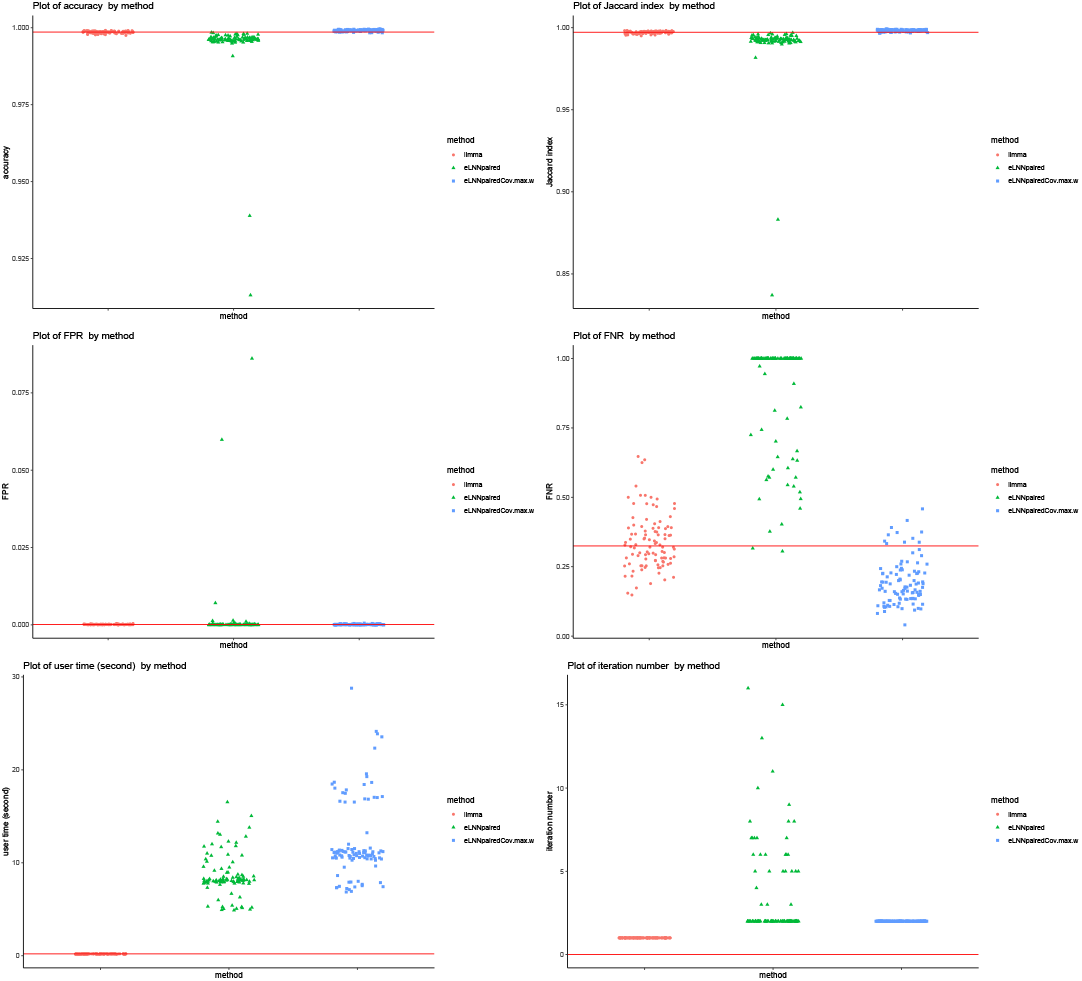
Jittered scatter plots of performance indices versus method for Scenario1 (number of pairs= 30). Red solid horizontal lines indicate the median performance indices of *limma*.

**Figure 4.**
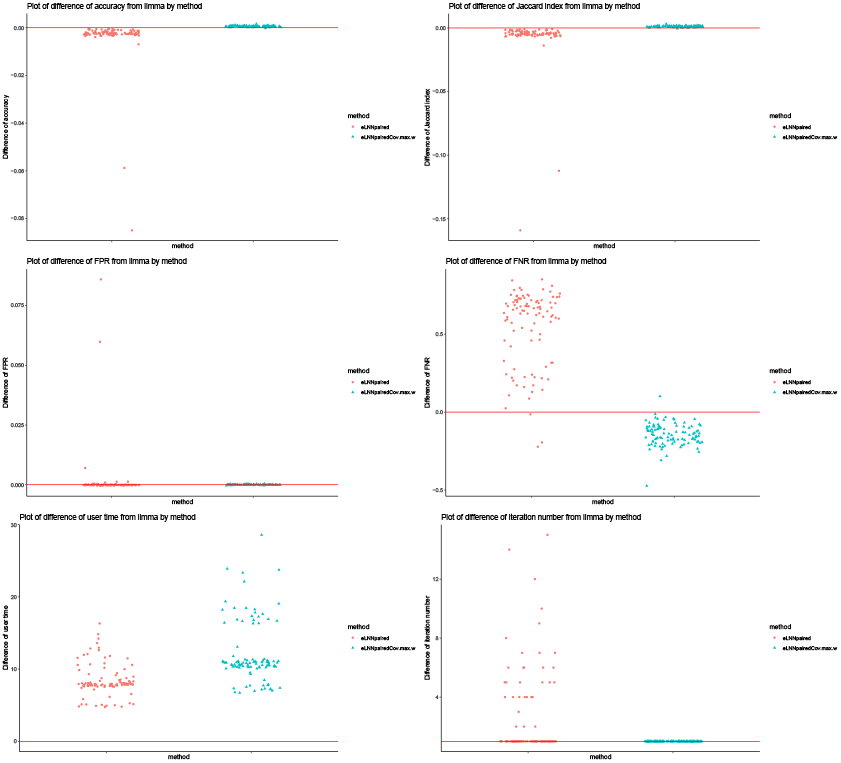
Jittered scatter plots of difference of performance indices versus method for Scenario1 (number of pairs= 30). Red solid horizontal lines indicate y-axis equal to zero, except for bottom-right plot, in which red sold horizontal line indicates y-axis equal to one.

The differences of performance indices are between *limma* and the other 2 methods (*eLNNpaired* and *eLNNpairedCov*). A positive difference indicates that the performance indices of the other 2 methods are larger than that of *limma*. A negative difference indicates that the performance indices of the other 2 methods are smaller than that of *limma*.

The upper panel of Figures 3 and 4 show that *eLNNpairedCov* has higher agreement indices (Jaccard and accuracy) than *limma*, which in turn has higher agreement indices than *eLNNpaired*.

The middle panel of Figures 3 and 4 show that the three methods have similar FPR, except that *eLNNpaired* has larger FPR than *eLNNpairedCov* and *limma* for a few simulated datasets. The middle panel also show that *eLNNpairedCov* has smaller FNR than *limma*, which in turn has smaller FNR than *eLNNpaired*.

The bottom panel of Figures 3 and 4 show that *limma* runs very fast, while both *eLNNpaired* and *eLNNpairedCov* run in reasonable time (i.e., less than 30 seconds per dataset that has *G* = 20, 000 genes and *n* = 30 subjects). On average *eLNNpairedCov* spends a little more time than *eLNNpaired*. The bottom panel of Figures 3 and 4 also show that *eLNNpairedCov*uses only 2 EM iterations, while *eLNNpaired* tends to use more EM iterations. Note that the EM iteration number for *limma* is set to be one, which does not use EM algorithm to obtain parameter estimates.

The simulation results for Scenario 2 (*n* = 100) are shown in Figures 6 and 7 in the Appendix, which have similar patterns to those for Scenario 1 (*n* = 30), except that both *eLNNpaired* and *eLNNpairedCov* have smaller FPR than *limma*. Note that *eLNNpairedCov* tends to have smaller FNR than *limma*, while *eLNNpaired* has huge FNR (close 1).

## 1 Discussion

In this article, we extended Li et al.’s (2017) mixture of Bayesian hierarchical models to allow the adjustment for potential confounding factors. This is novel in that to the best of our knowledge, all existing clustering methods do not yet have the capacity to adjust for covariates.

The proposed approach is different from transcript-wise test with controlling multiple testing in that it does not involve hypothesis testing. Hence, no multiple test controlling is needed. The simulation study showed that if the difference of gene expression between samples before treatment and samples after treatment follows the mixture of Bayesian hierarchical models in Section, then the proposed method can outperform *limma*, which currently is a fast and powerful transcript-wise test method. The real data analysis also showed the proposed method *eLNNpairedCov* can detect more differentially expressed gene transcripts, which include the transcripts detected by *limma*.

Although we classify genes to three distinct clusters, the transitions between these clusters could be smooth. This would be reflected by a gene’s posterior probability that might be large in two of three clusters, e.g., 0.49 for cluster 1, 0.01 for cluster 2, and 0.5 for cluster 3. On the other hand, expression changes could be split up into more than 3 clusters, e.g., groups behaving differently. In this article, we are only interested in identifying three clusters of genes: over-expressed in condition 1, under-expressed in condition 1, and non-differentially expressed.

There are other model-based clustering methods in literature, such as^16^. However, they were not designed to detect differentially expressed genes. For example, we can set the number *K* of clusters as 3 for their model. However, there is no constraints that the intercepts for the three clusters have to be positive, negative, and zero. That is, the three clusters identified might not correspond to over-expressed, under-expressed, and non-differentially expressed genes.

RNAseq and single-cell RNAseq data are cutting-edge tools to investigate molecular mechanisms of complex human diseases. However, it is quite challenging to analyze these count data with inflated zero counts. In future, we will evaluate if eLNNpairedCov can be used to analyze single-cell RNAseq data by first transforming counts to continuous scale (e.g., via VOOM^9^ or countTransformers^10^) and then to apply eLNNpairedCov to the transformed data.

We implemented the proposed methods to an R package *eLNNpairedCov*, which will be freely available to researchers.

## Author contributions statement

Conceptualization, W.L. and W.Q.; methodology, Y.Z., W.L, and W.Q.; software, Y.Z. and W.Q.; validation, Y.Z., W.L. and W.Q.; formal analysis, Y.Z. and W.Q.; investigation, W.L.; resources, W.L.; data curation, Y.Z.; writing—original draft preparation, Y.Z.; writing—review and editing, W.L. and W.Q.; visualization, Y.Z. and W.Q.; supervision, W.L. and W.Q.; project administration, W.L.; funding acquisition, W.L. All authors have read and agreed to the published version of the manuscript.

## Data availability statement

The data presented in this study are openly available in [https://www.ncbi.nlm.nih.gov/geo/query/acc.cgi?acc=GSE24742]

## Additional information

The authors declare no potential conflict of interests.

## Appendix

### Histogram of estimated standard deviations of log2 differences of within-subject gene expression for GSE24742

**Figure 5.**
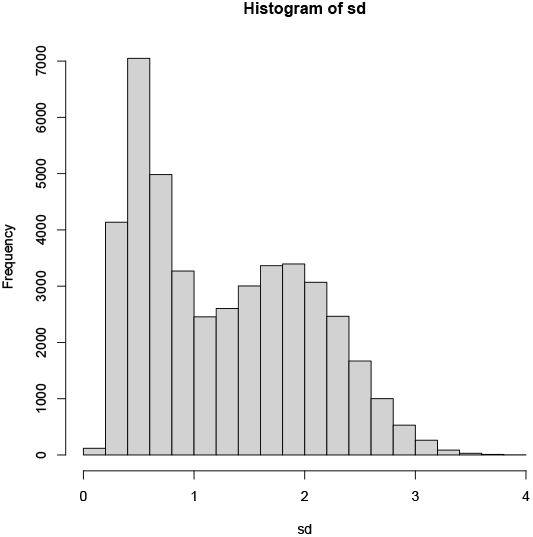
Histogram of the estimated standard deviations of log2 differences of within-subject gene expression for 43,506 gene probes in GSE24742.

#### Additional results

**Figure 6.**
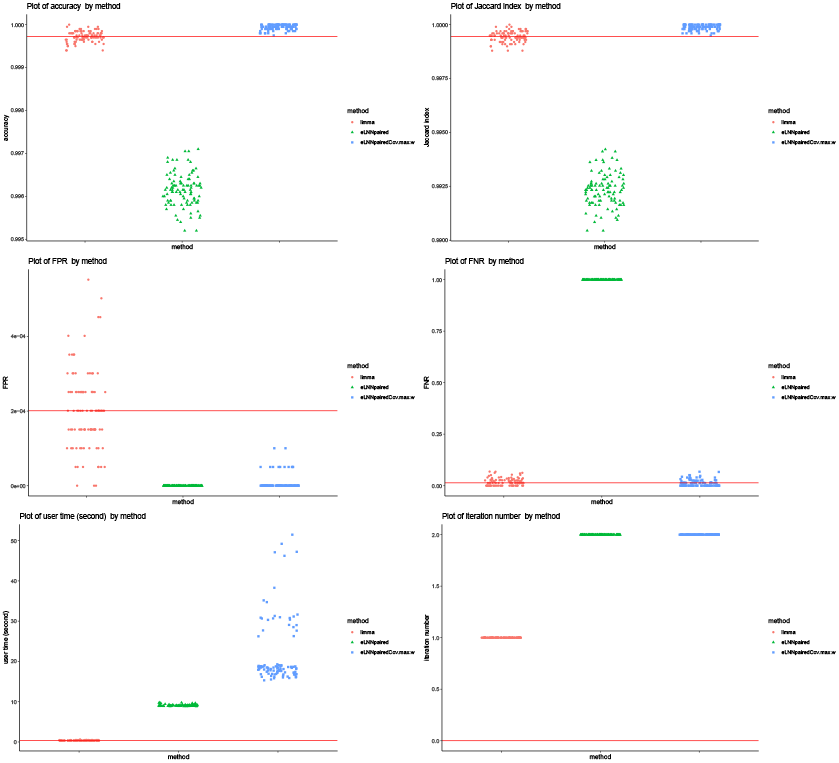
Jittered scatter plots of performance indices versus method for Scenario2 (number of pairs= 100). Red solid horizontal lines indicate the median performance indices of *limma*.

**Figure 7.**
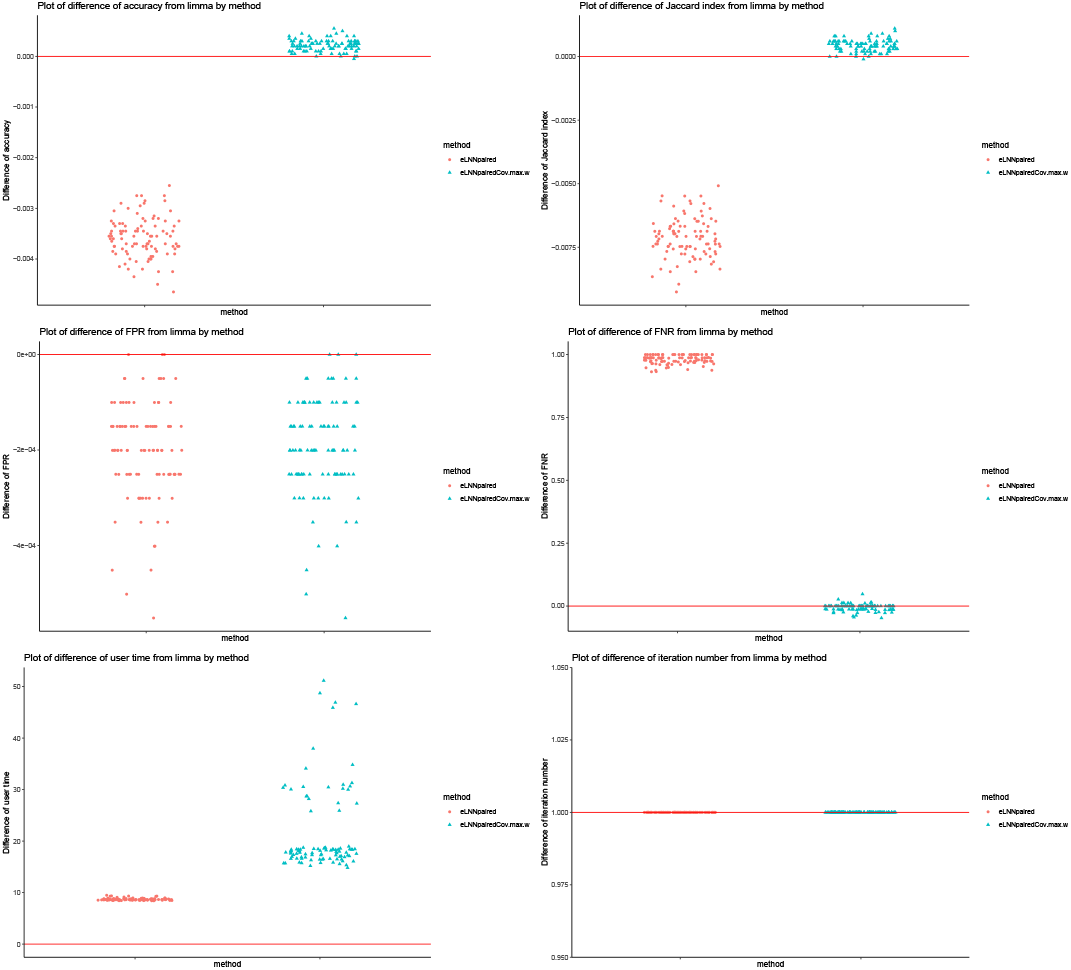
Jittered scatter plots of difference of performance indices versus method for Scenario2 (number of pairs= 100). Red solid horizontal lines indicate y-axis equal to zero, except for bottom-right plot, in which red sold horizontal line indicates y-axis equal to one.

#### Marginal density functions

The marginal density function of **d**_*g*_is :

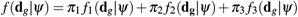

where *f*_1_(**d**_*g*_| *ψ*), *f*_2_(**d**_*g*_ |*ψ*), *f*_3_(**d**_*g*_ |*ψ*) are the marginal densities of the 3 clusters, and *π* = (*π*_1_, *π*_2_, *π*_3_) is the cluster proportion.

Then we have the marginal distribution for each cluster:

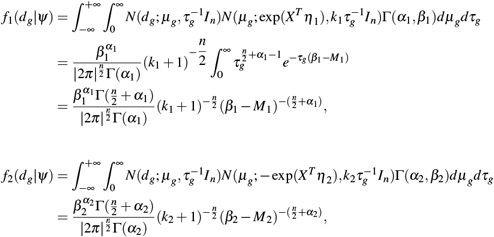

where *X*^*T*^ is the *n* × (*p* + 1) design matrix.

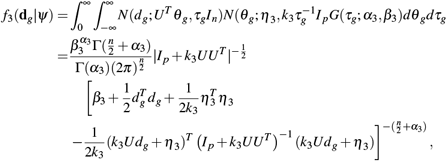

where *U*^*T*^ is the *n × p* design matrix without intercept column.

Here

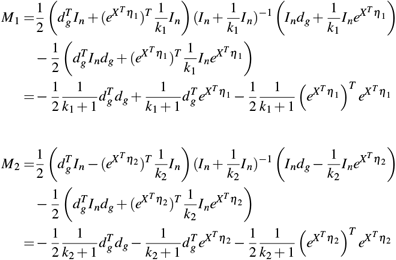

Note that each marginal density is the density of a multivariate t distribution with special structures of mean vector and covariance matrix.

#### Log marginal densities

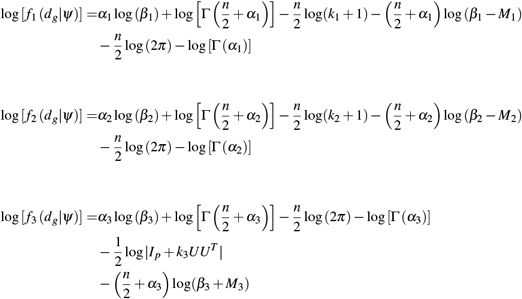

Here

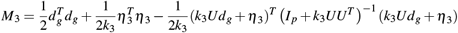

#### EM algorithm

Let *z*_*g*_ = (*z*_*g*1_, *z*_*g*2_, *z*_*g*3_) to be the indicator vector indicating if gene transcript *g* belongs to a cluster or not. To stablize the estimate of *π* when *π*_*c*_ is very small, we assume that the cluster mixture proportions *π* follows a symmetric Dirichlet *D*(**b**) distribution, i.e., 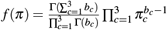. Therefore, the likelihood function for the complete data (**d, z**, *π*) is

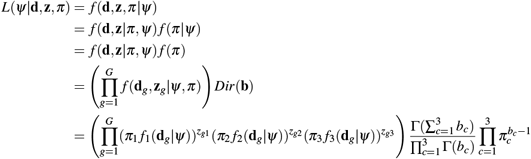

Then the log complete-data likelihood function is:

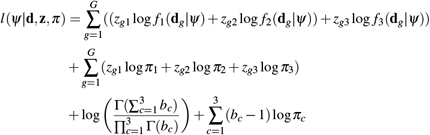

The EM algorithm is used to estimate parameters *π* and *ψ*. Since *z* is unknown random vector, we integrate it out from the log complete-data likelhood function. Here, **z**_*g*_ = (*z*_*g*1_, *z*_*g*2_, *z*_*g*3_).

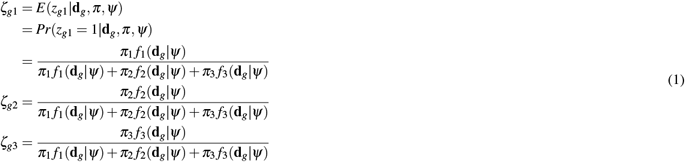

#### E-step

Denote *Q*^(*t*)^ (*π, ψ*|*d, z*^(*t*)^, *π*^(*t*)^)as the expected log complete-data likelihood function at *t*-th iteration of the EM algorithm, we have

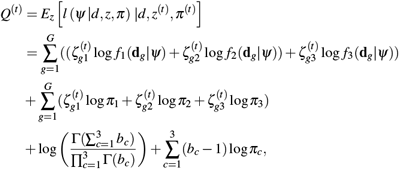

where

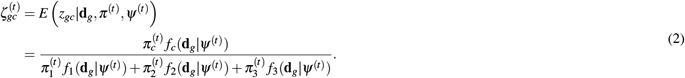

#### M-step

Maximize *Q*^(*t*)^ (*π, ψ*|*d, z*^(*t*)^, *π*^(*t*)^) to find the optimal values of *π* and *ψ*, and use these optimal values as estimates for the parameters *π* and *ψ*.

To maximize *Q*^(*t*)^ (*π, ψ*|*d, z*^(*t*)^, *π*^(*t*)^), we use the “L-BFGS-B” method developed by Byrd et al. (1995)^17^, which utilizes the first partial derivatives of *Q*^(*t*)^ (*π, ψ*|*d, z*^(*t*)^, *π*^(*t*)^) and allows box constraints, that is each variable can be given a lower and/or upper bound.

#### First Derivatives

For *π*_1_, *π*_2_, *π*_3_, we have:

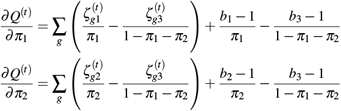

Since 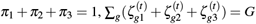, and letting *b*_1_ = *b*_2_ = *b*_3_ = 2, *∂Q*^(*t*)^*/∂π*_1_ = *∂Q*^(*t*)^*/∂π*_2_ = 0, we have:

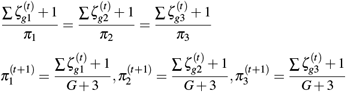

##### For OE genes

For *α*_1_, *β*_1_, *k*_1_ and *η*_1_, we have

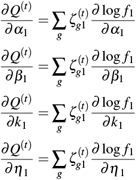

Now, we would like to take partial derivatives of log *f*_1_ with respect to *α*_1_, *β*_1_, *k*_1_ and *η*_1_. And we have:

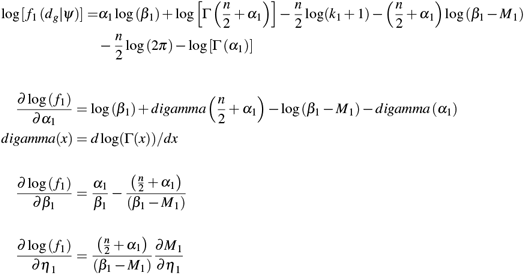

Denote

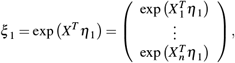

where

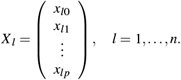

Then we can rewrite *M*_1_ as

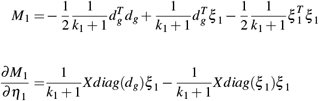

As for *k*_1_, we have

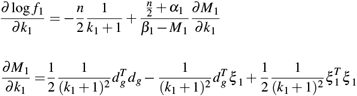

##### For UE gene probes

For *α*_2_, *β*_2_, *k*_2_ and *η*_2_, we have

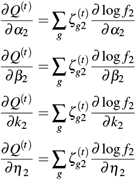

Take partial derivatives of log *f*_2_ with respect to *α*_2_, *β*_2_, *k*_2_ and *η*_2_. And we have:

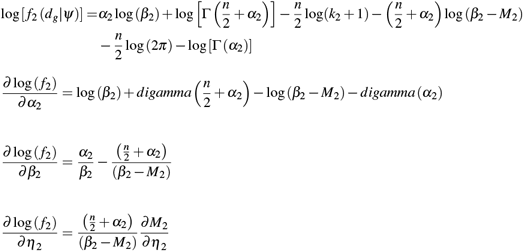

Denote

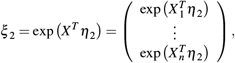

where

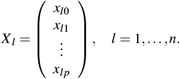

Then we can rewrite *M*_2_ as

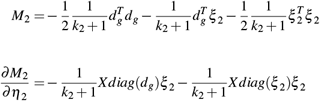

For *k*_2_, we have

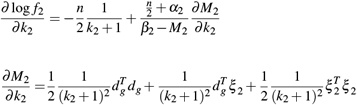

##### For NE gene probes

For *α*_3_, *β*_3_, we have

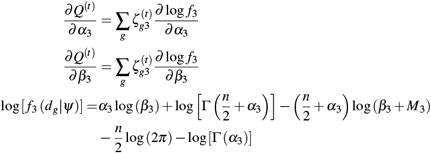

Here

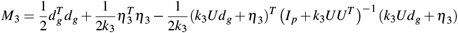

Therefore, take the derivatives of log *f*_3_, we have

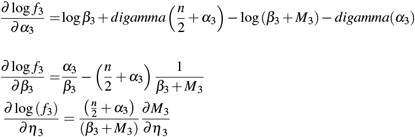

For 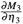, we consider the following two components,

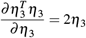

And we have (Matrix Cookbook^18^, equation(85)), given **W** is symmetric,

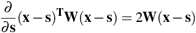

Then,

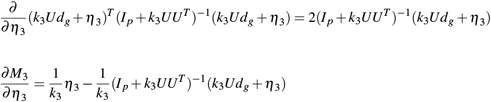

For *k*_3_, we have

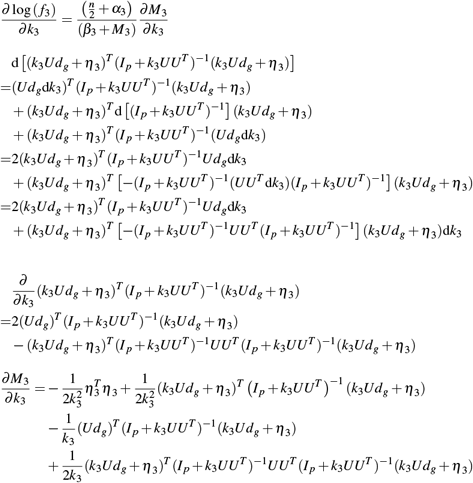

#### Initial parameter estimates

We first run *limma* to get false discovery rate (FDR) adjusted p-value of moderated t-test for each gene transcript. We then partition the gene transcripts to 3 clusters based on the FDR-adjusted p-values and signs of the test statistics: (1) gene transcripts with positive test statistics and FDR-adjusted p-values *<* 0.05 (denoted as OE); (2) gene transcripts with negative test statistics and FDR-adjusted p-values *<* 0.05 (denoted as UE); and (3) the remaining gene transcripts (denoted as NE). If numbers of transcripts in OE and/or UE are smaller than 5, we then use 100 transcripts having the smallest un-adjusted p-values to find initial OE and UE transcripts. We next estimate the cluster mixture proportions (*π*_1_, *π*_2_, *π*_3_) by the 3 cluster sizes dividing the total number of gene transcripts.

Within each cluster, we use moment estimator to get the initial estimates of model parameters. In our case, we use sample median (med) and sample median absolute deviation (mad) to get robust estimates of mean and standard deviation, based on which we get the initial estimate of *ψ*.

##### OE gene transcripts

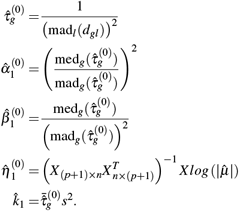

where

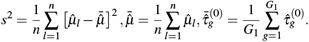

##### UE gene transcripts

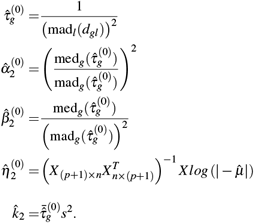

where

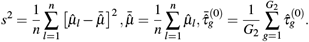

Here, we use 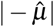 to make sure it is positive.

##### NE gene transcripts

The initial values are

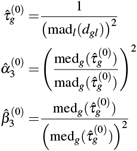

As for *η*_3_, we first need to get 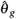 With linear regression, we have

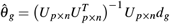

Then we can get

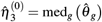

And

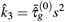

where

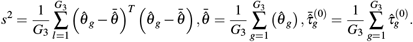

#### Bounds for hyper-parameters

Although we can re-parameterize hyper-parameters to make sure *α*_*c*_ > 0, *β*_*c*_ > 0, 0 *< k*_*c*_ *<* 1, *c* = 1, 2, 3, and to transform constrained optimization problem to un-constrained optimization problem, the results are usually not good due to non-linear optimization in the M-step of the EM algorithm. To avoid un-expected large values of estimated *α*_*c*_, *β*_*c*_, and/or *k*_*c*_ in numerical optimization, we set constraints 0.001 *< α*_*c*_ *<* 6, 0.001 *< β*_*c*_ *<* 6, 0.001 *< k*_*c*_ *<* 0.9999.

If initial estimates 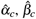, or 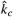 are outside their ranges, then we randomly choose a number between the range as the initial parameter estimates.

In the M-step of the EM iterations, we also require each element of *η*_*c*_, *c* = 1, 2, 3, be within the interval [−10, 10].

If at least one element of *ψ*^(*t*)^ are outside their ranges in the *t*-th iteration of the EM algorithm, we then set final estimate of *ψ* as *ψ*^(*t*−1)^, and then only update 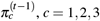, until convergence criterion satisfied.

The M-step in the EM algorithm is to maximize *Q* function, which is a non-linear function of model parameters *π*_1_, *π*_2_, *π*_3_, *ψ*. It is difficult to optimize non-linear functions. The above constraints and procedure produced good results in the real and simulation data analyses in this article.

#### Information for differentially expressed transcripts for GSE24742

The information about the 6 UE transcripts detected by *limma* is shown in Table 1.

**Table 1.** Transcript IDs, gene symbols, chromosome numbers of the 6 UE transcripts detected by *limma*

| ID | gene | chr |
| --- | --- | --- |
| 1560920_s_at | LOC102725382 | 9 |
| 202067_s_at | LDLR | 19 |
| 207306_at | TCF15 | 20 |
| 219249_s_at | FKBP10 | 17 |
| 222419_x_at | UBE2H | 7 |
| 224507_s_at | MGC12916 | 17 |

The information about the 9 OE transcripts detected by *eLNNpairedCov* is shown in Table 2.

**Table 2.** Transcript IDs, gene symbols, chromosome numbers of the 9 OE transcripts detected by *eLNNpairedCov*

| ID | gene | chr |
| --- | --- | --- |
| 1552808_at | ADAMTSL1 | 9 |
| 1570121_at | ZNF365 | 10 |
| 222223_s_at | IL36RN | 2 |
| 231595_at | MRVI1-AS1 | 11 |
| 232186_at | FITM2 | 20 |
| 233466_at | LOC101928729 | 17 |
| 234138_at | LOC105378749 | 1 |
| 238249_at | LOC101927499 | 22 |
| 244557_at | LINC00610 | 11 |

The information about the 83 UE transcripts detected by *eLNNpairedCov* is shown in Table 3.

**Table 3.**
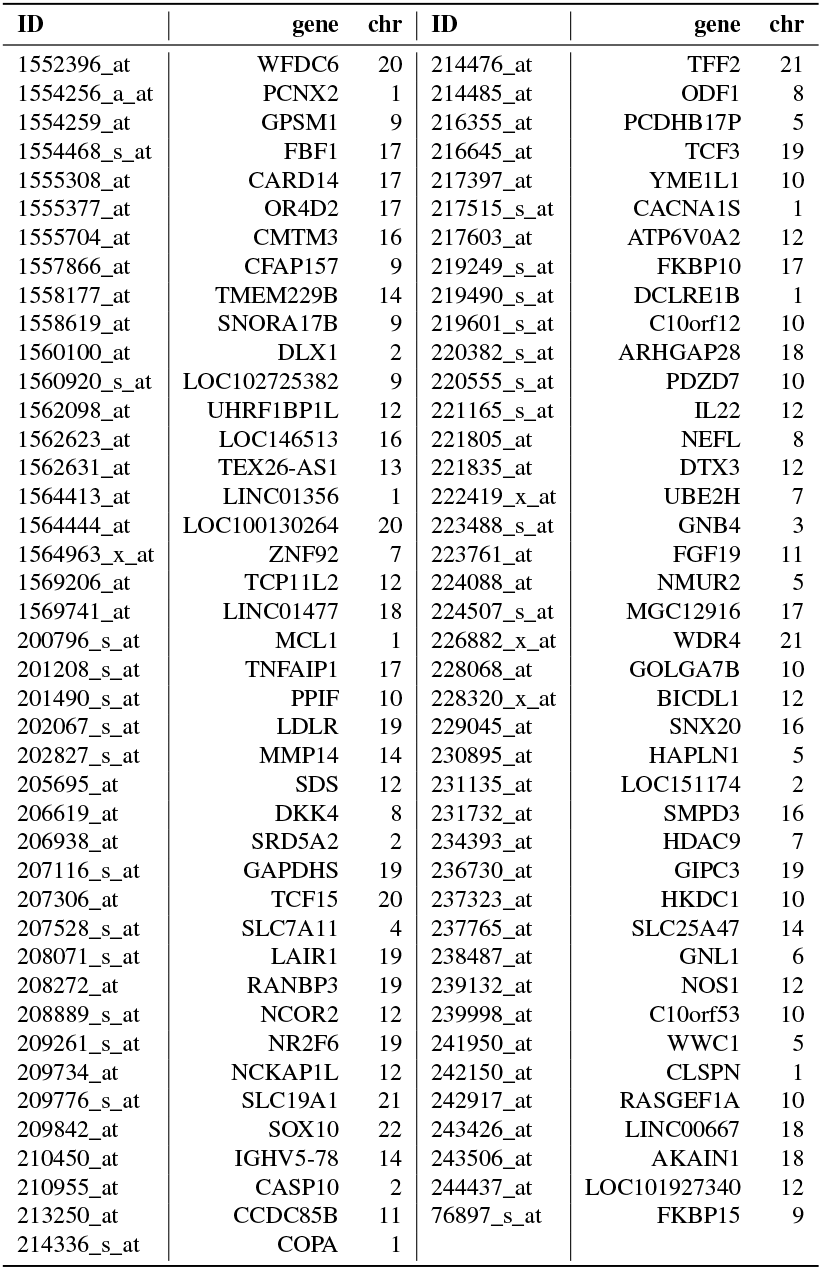
Transcript IDs, gene symbols, chromosome numbers of the 83 UE transcripts detected by *eLNNpairedCov*

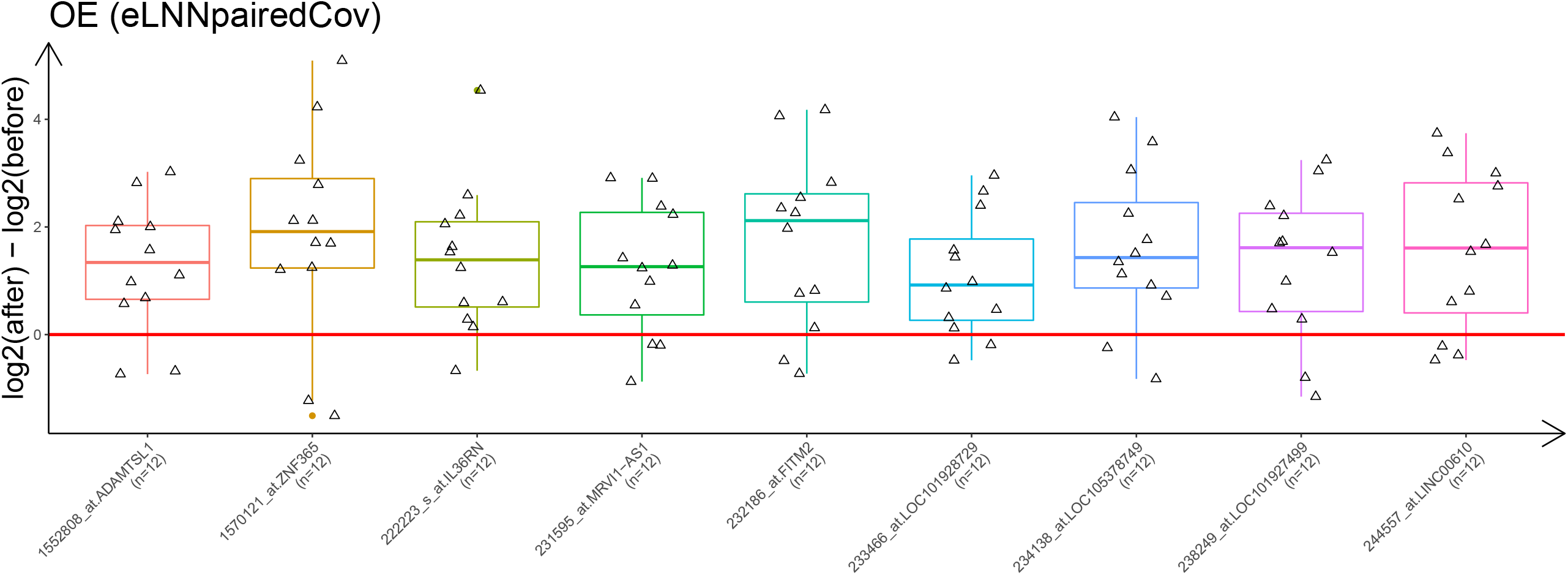

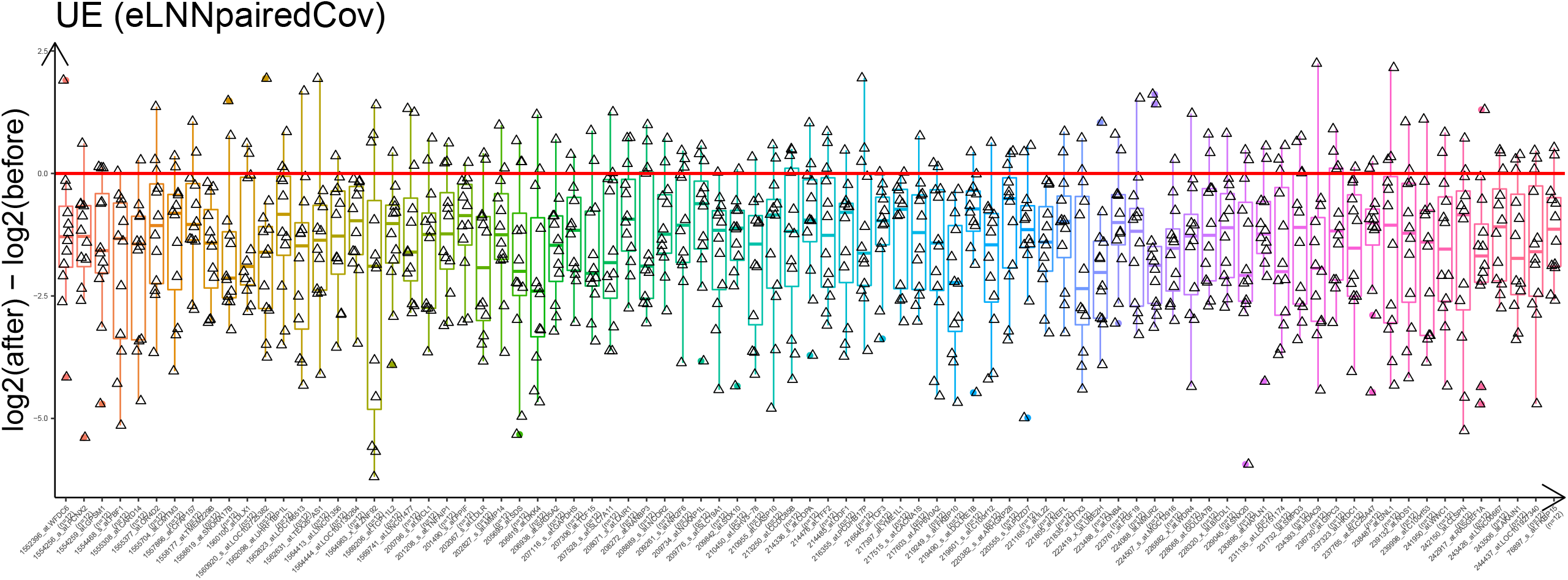

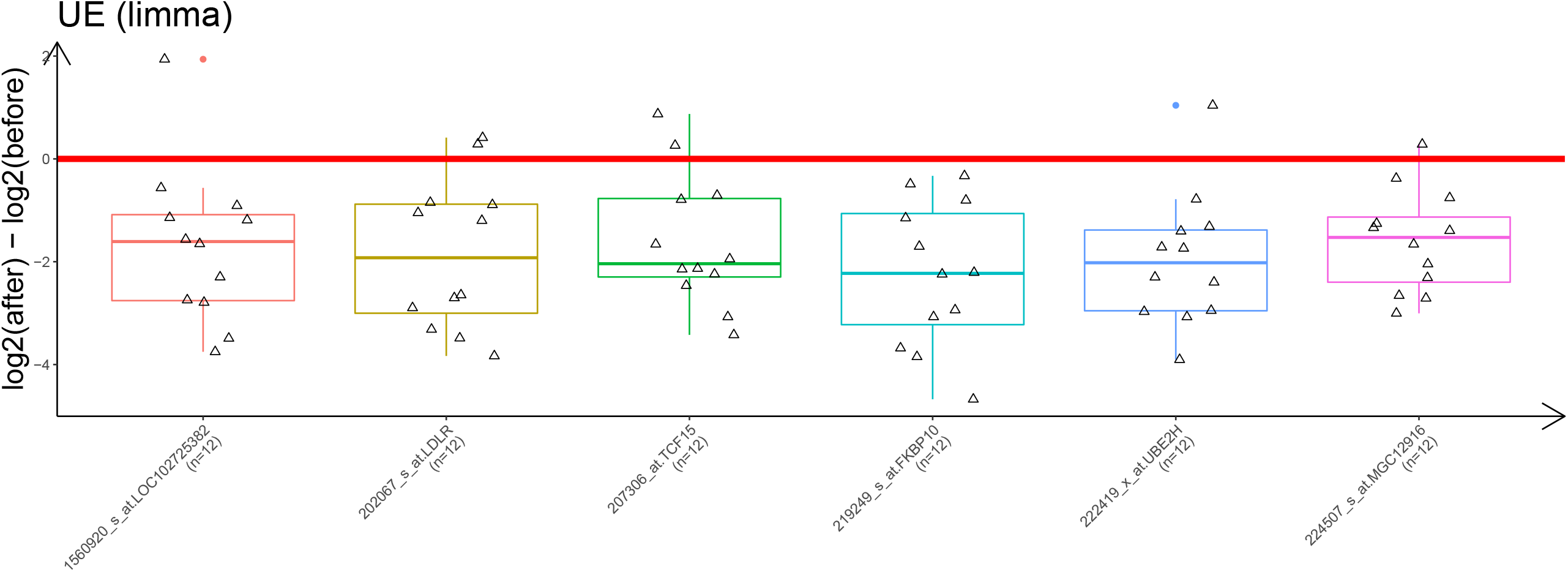

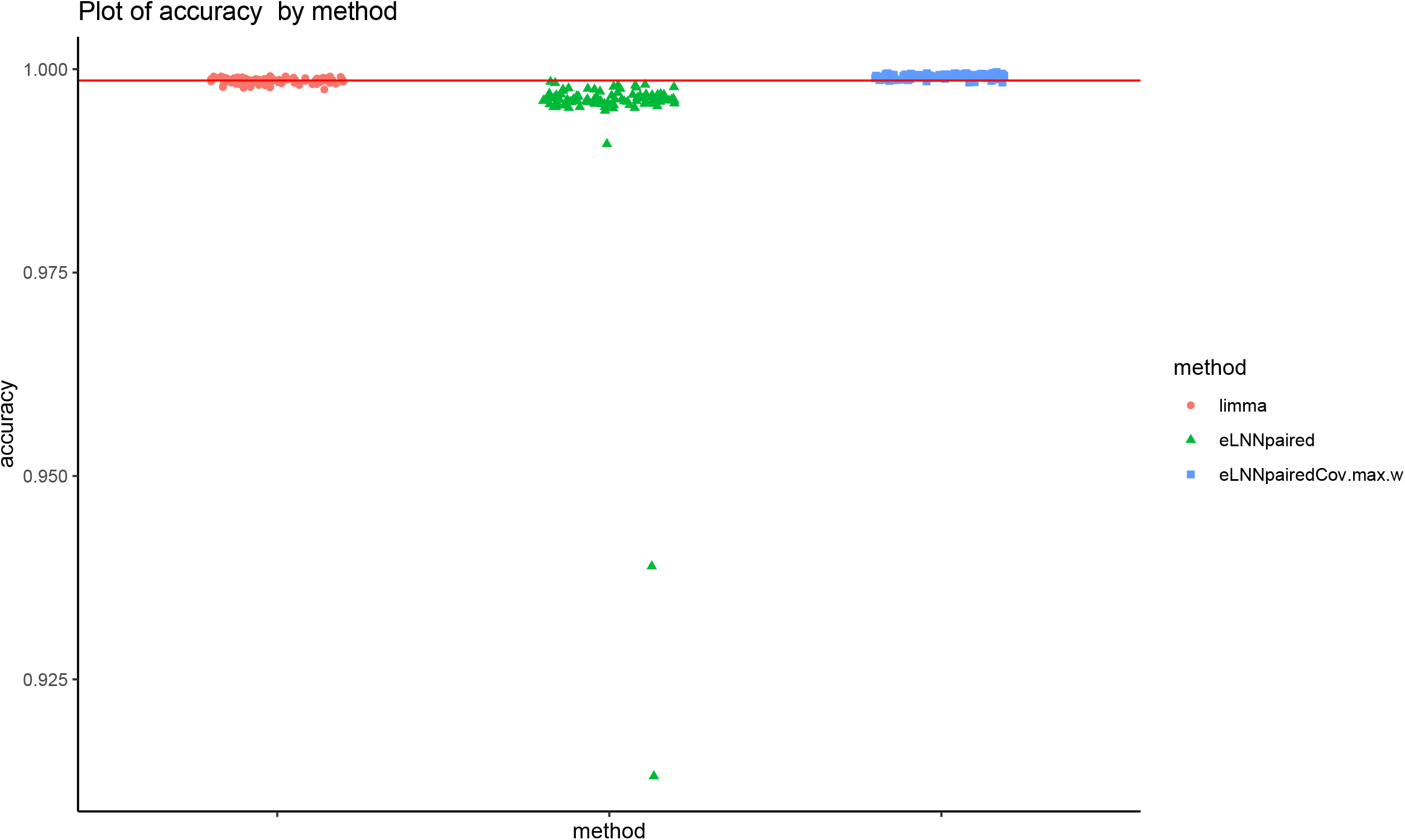

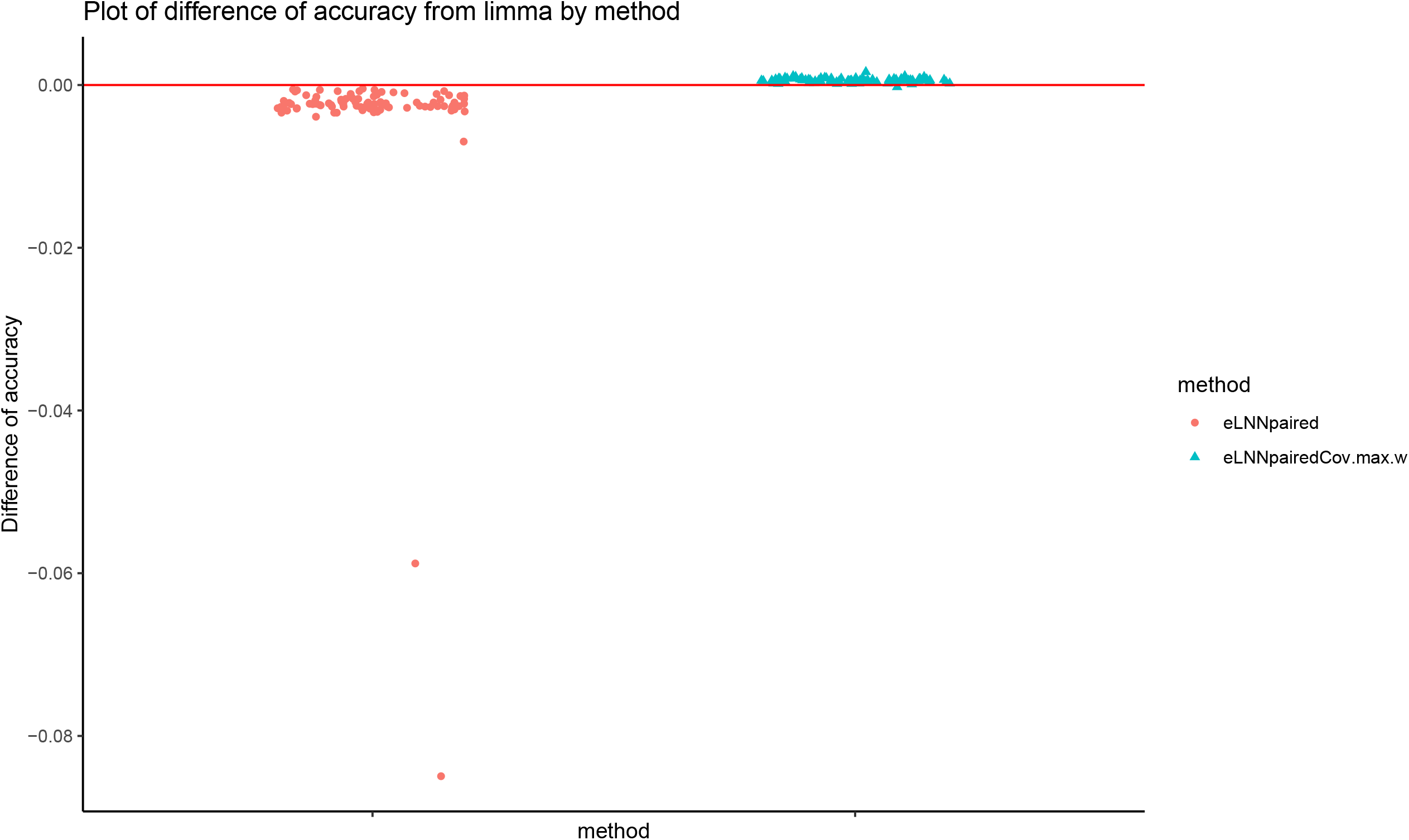

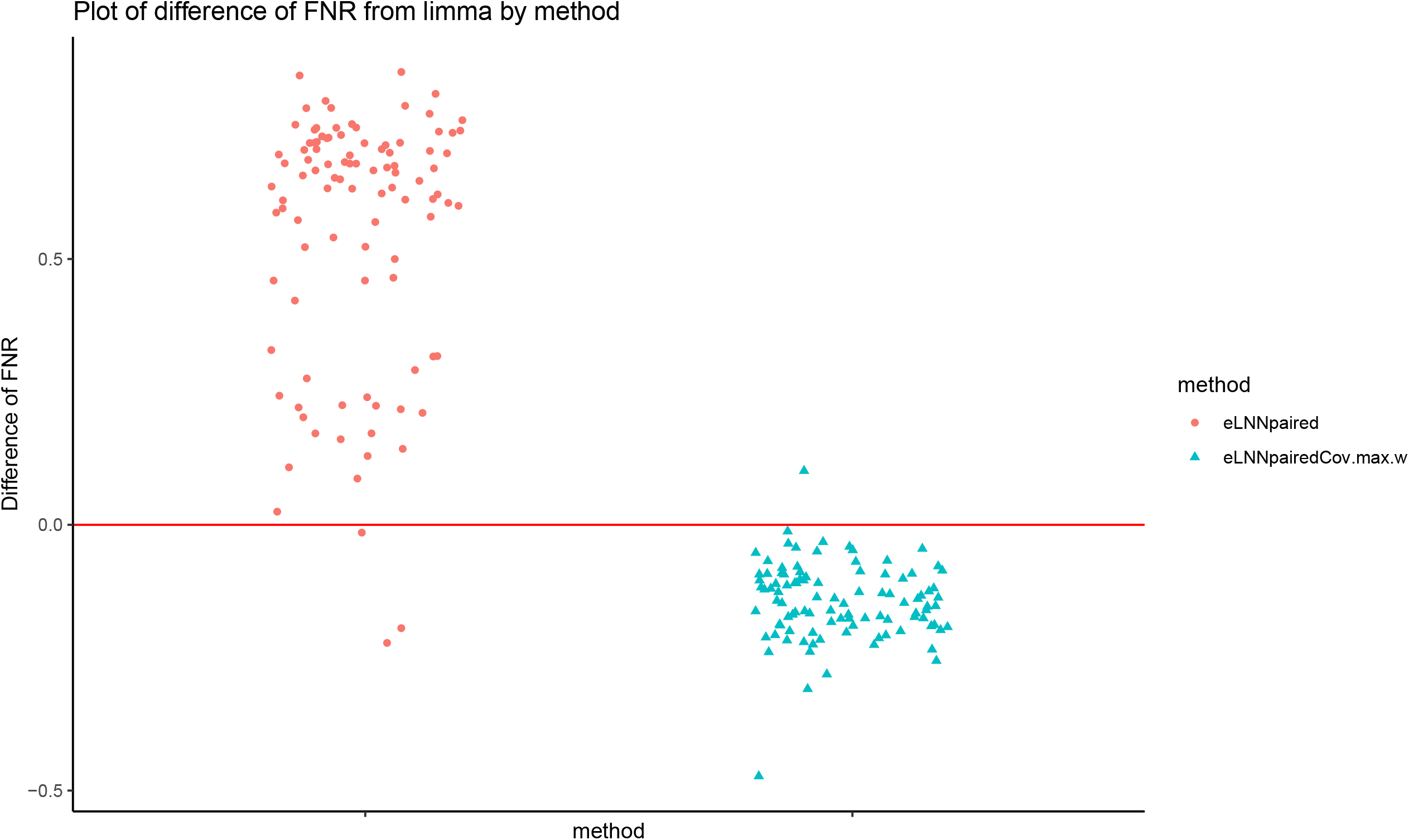

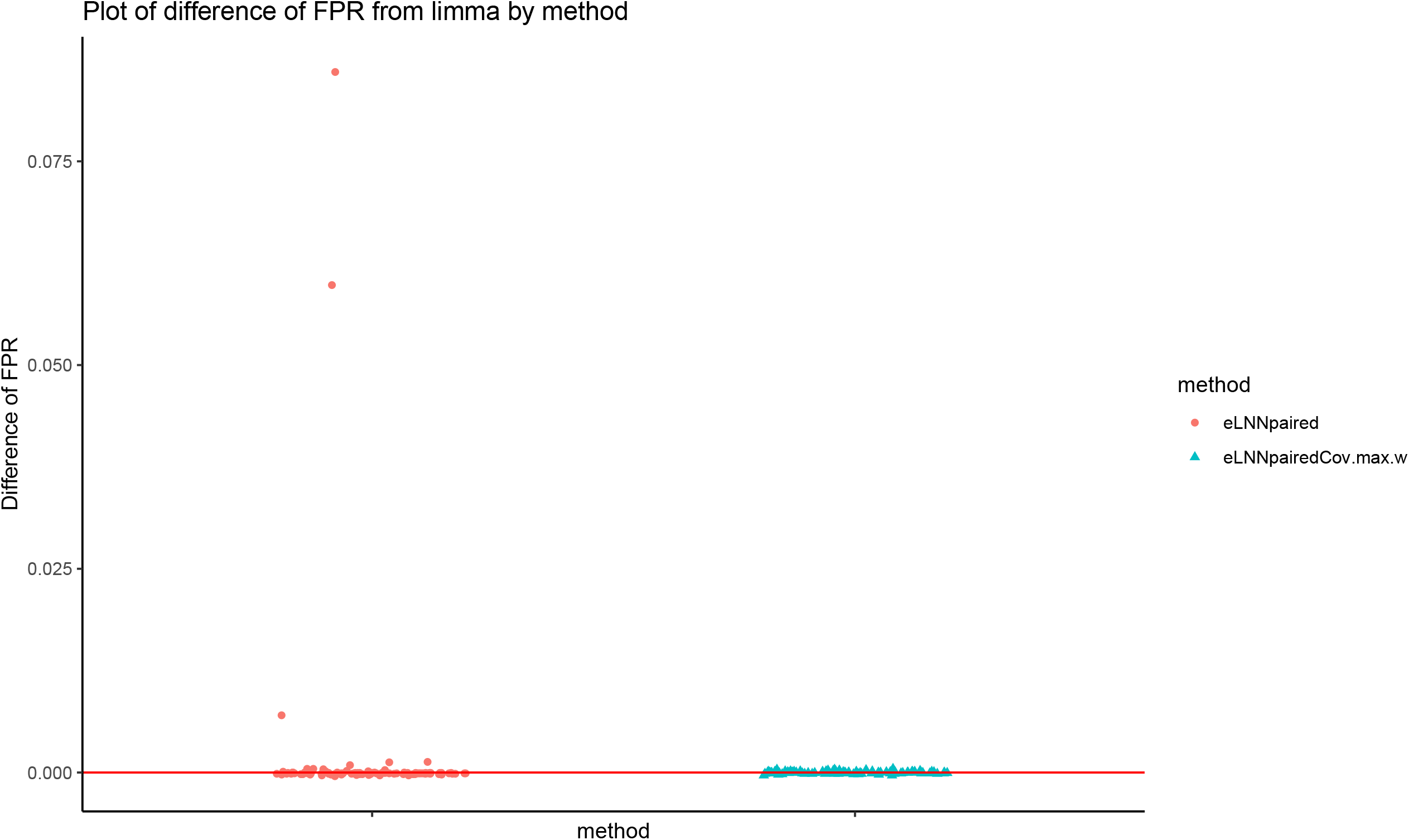

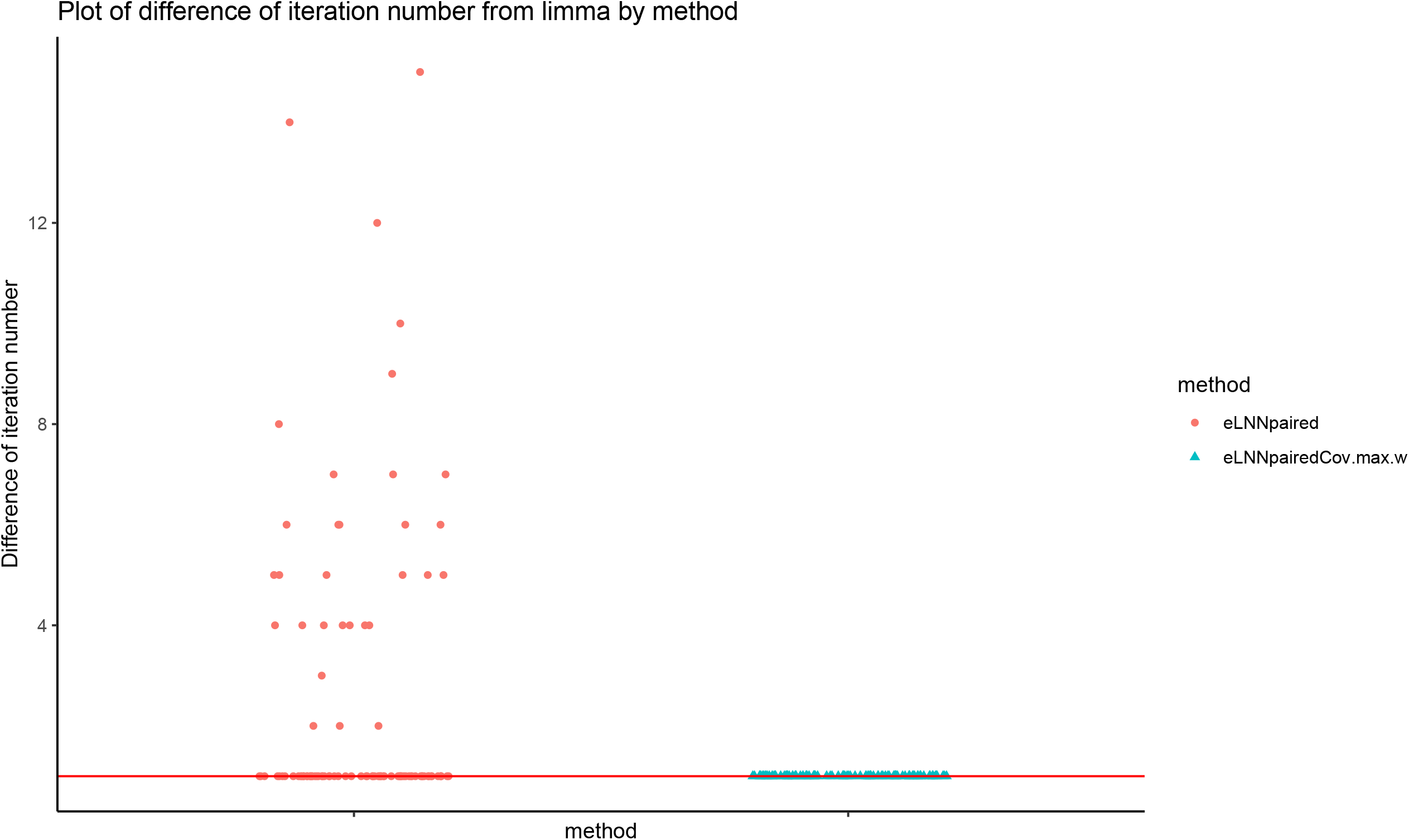

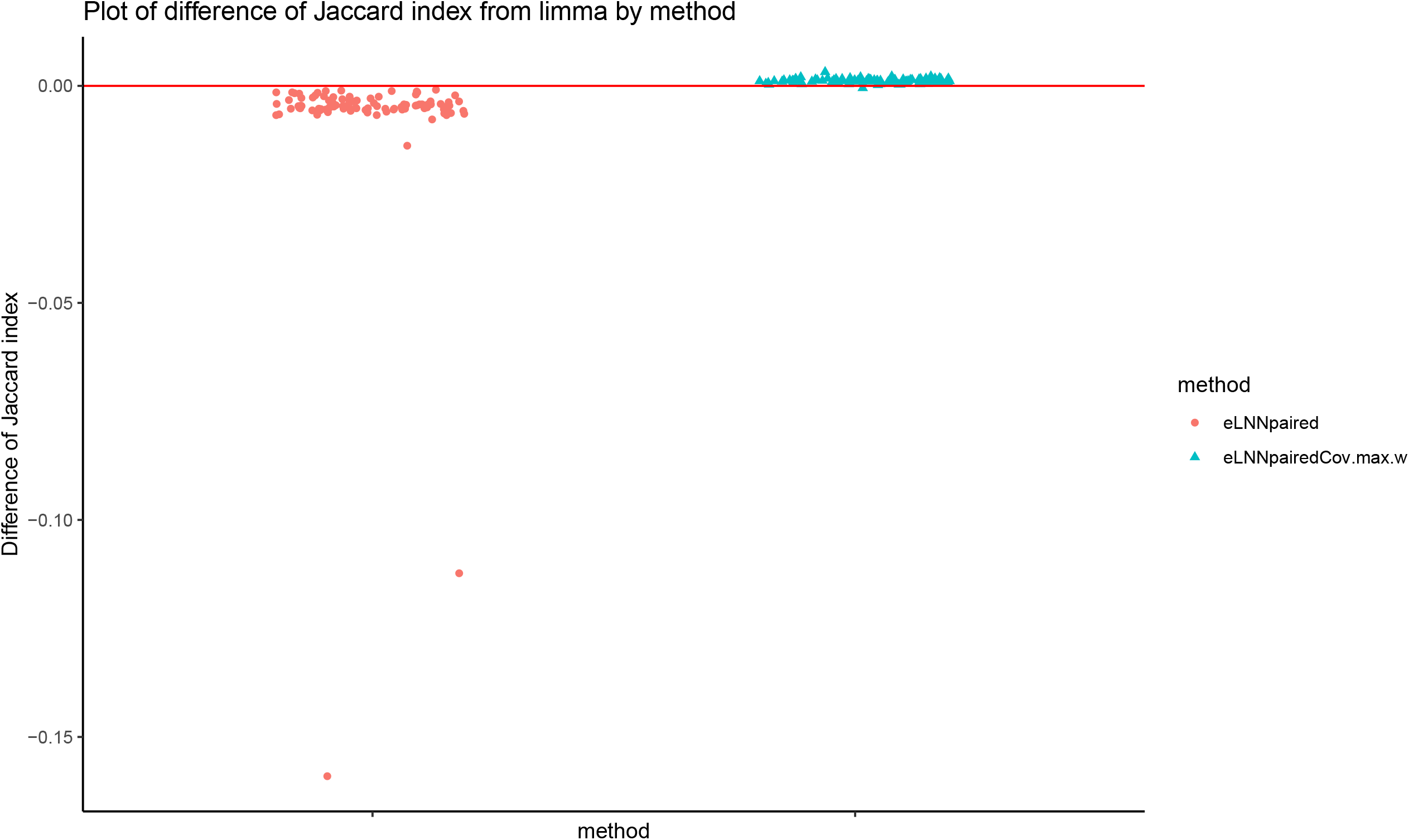

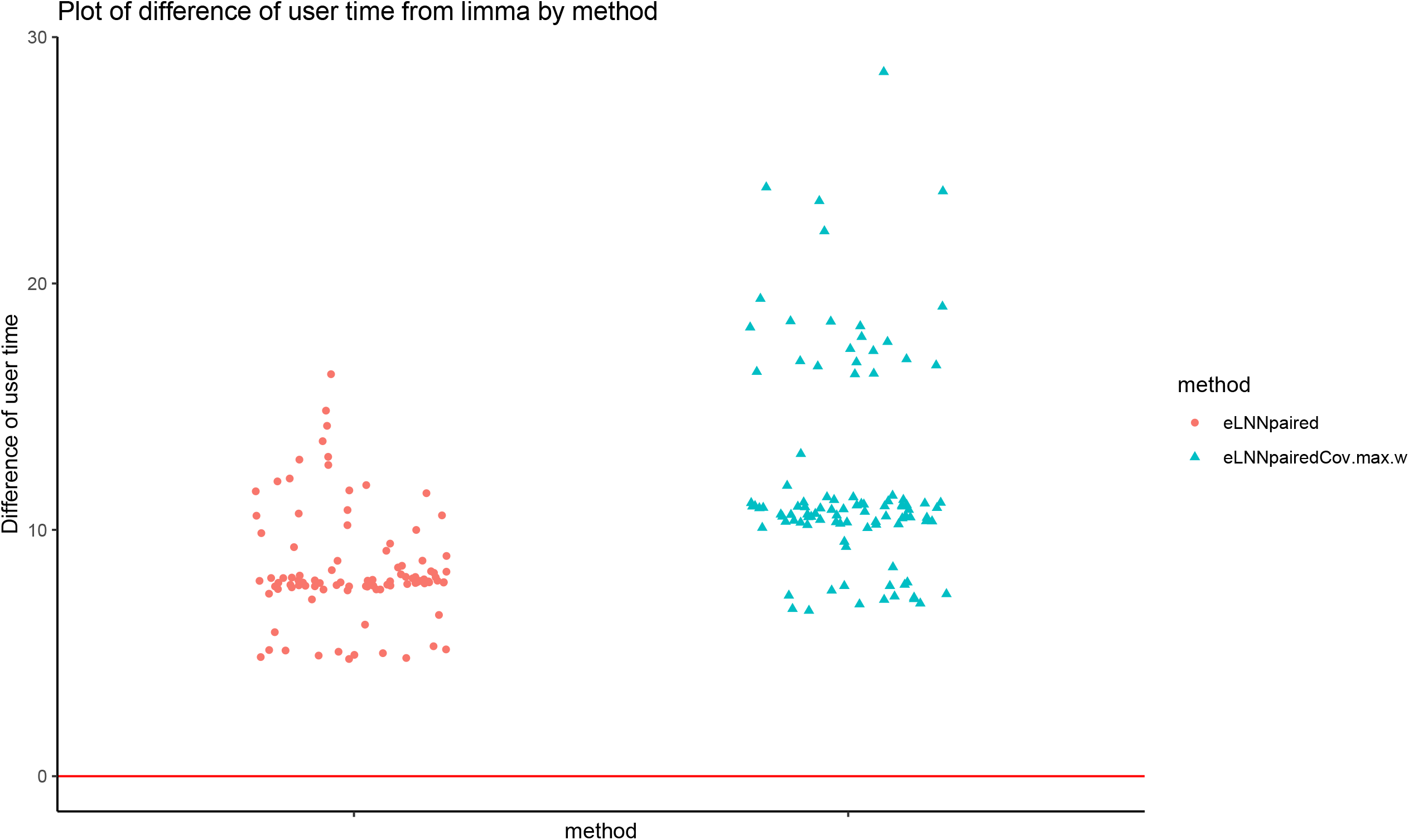

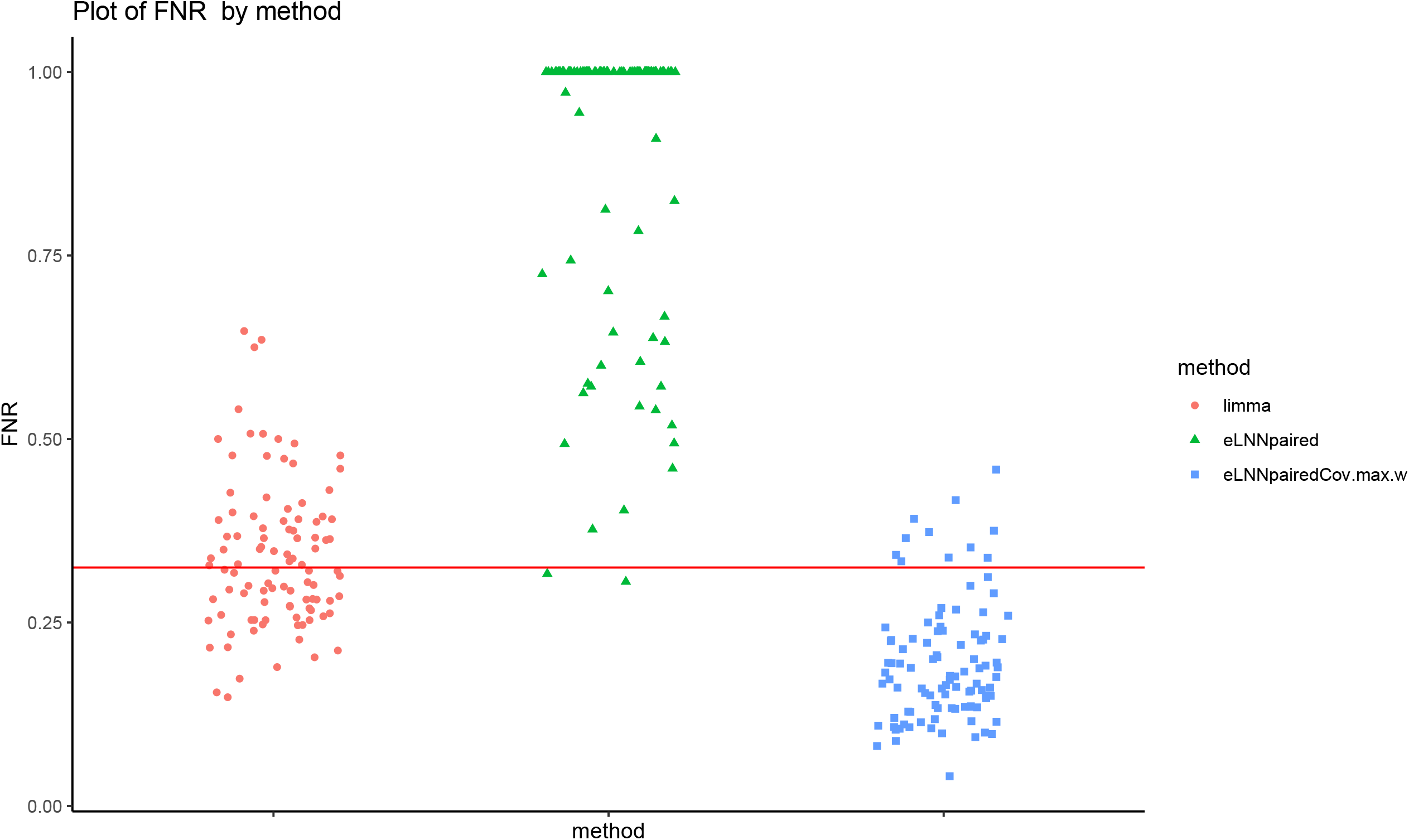

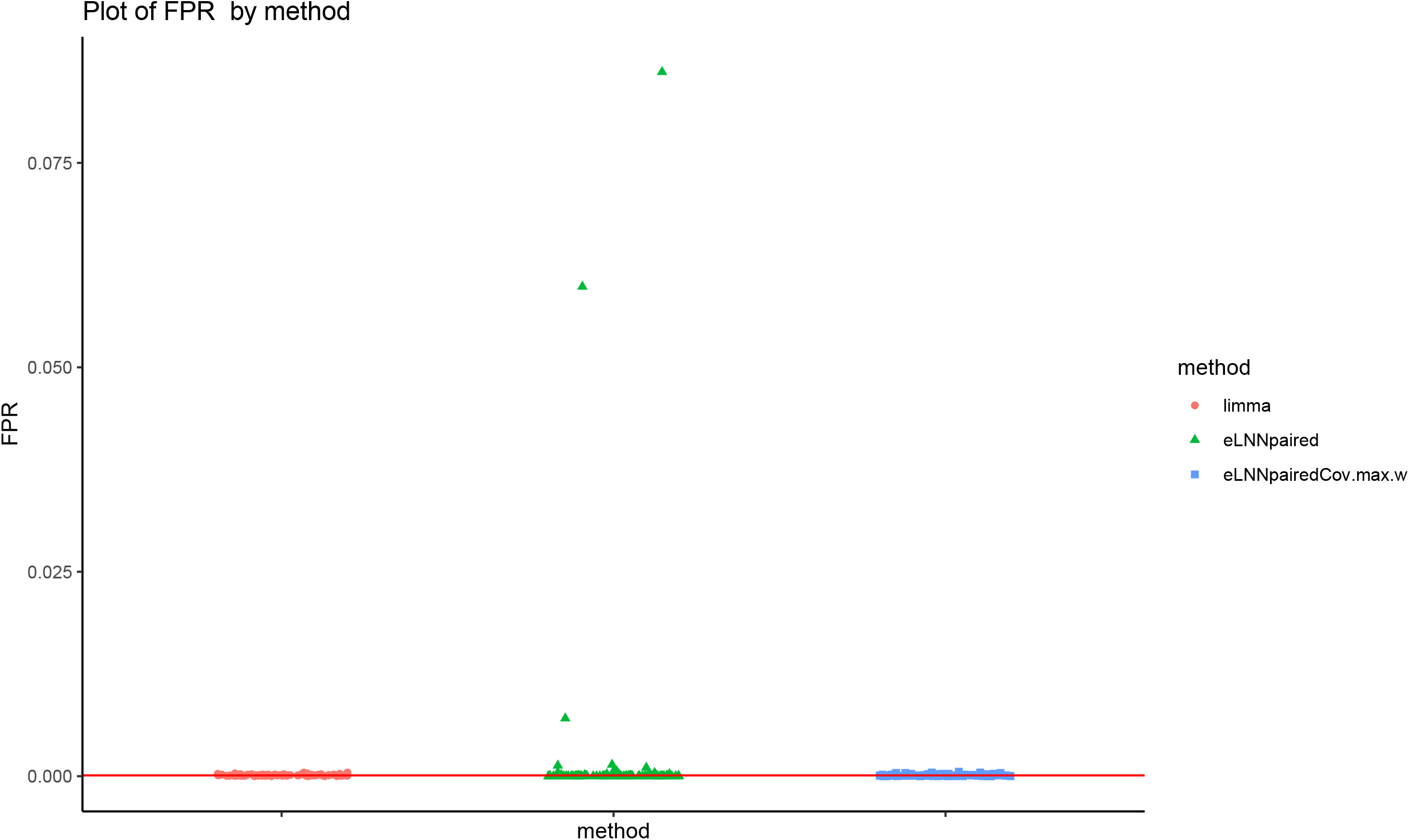

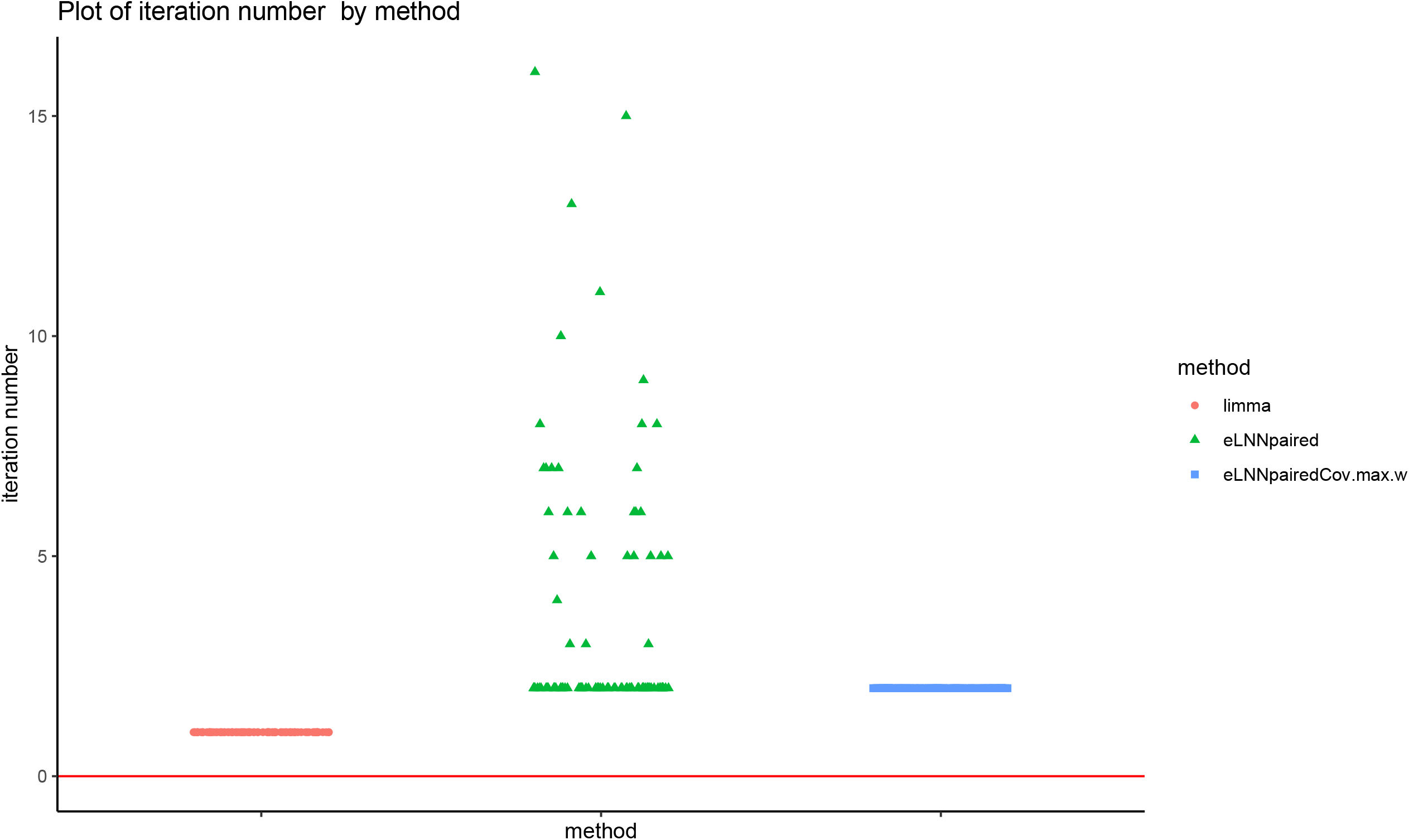

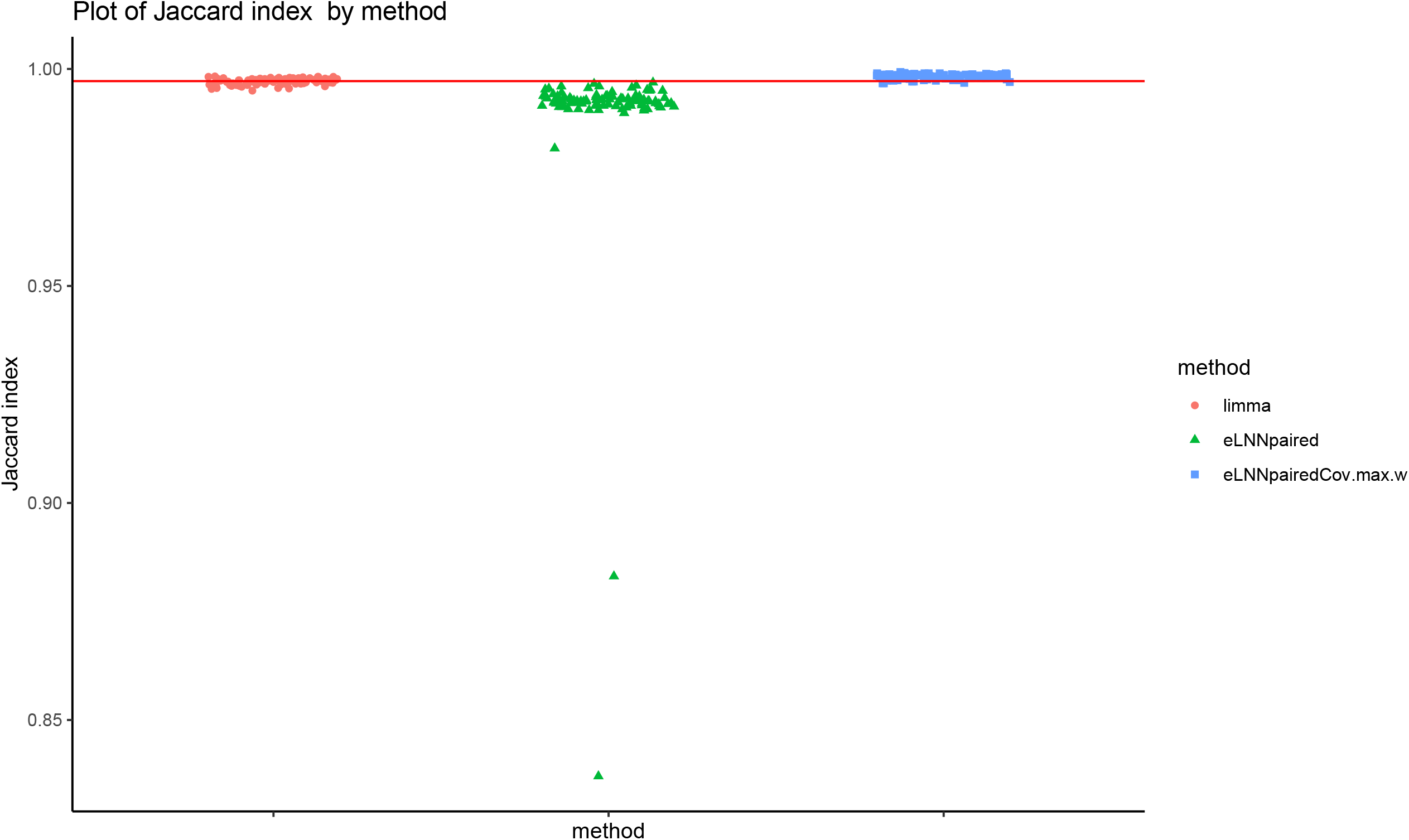

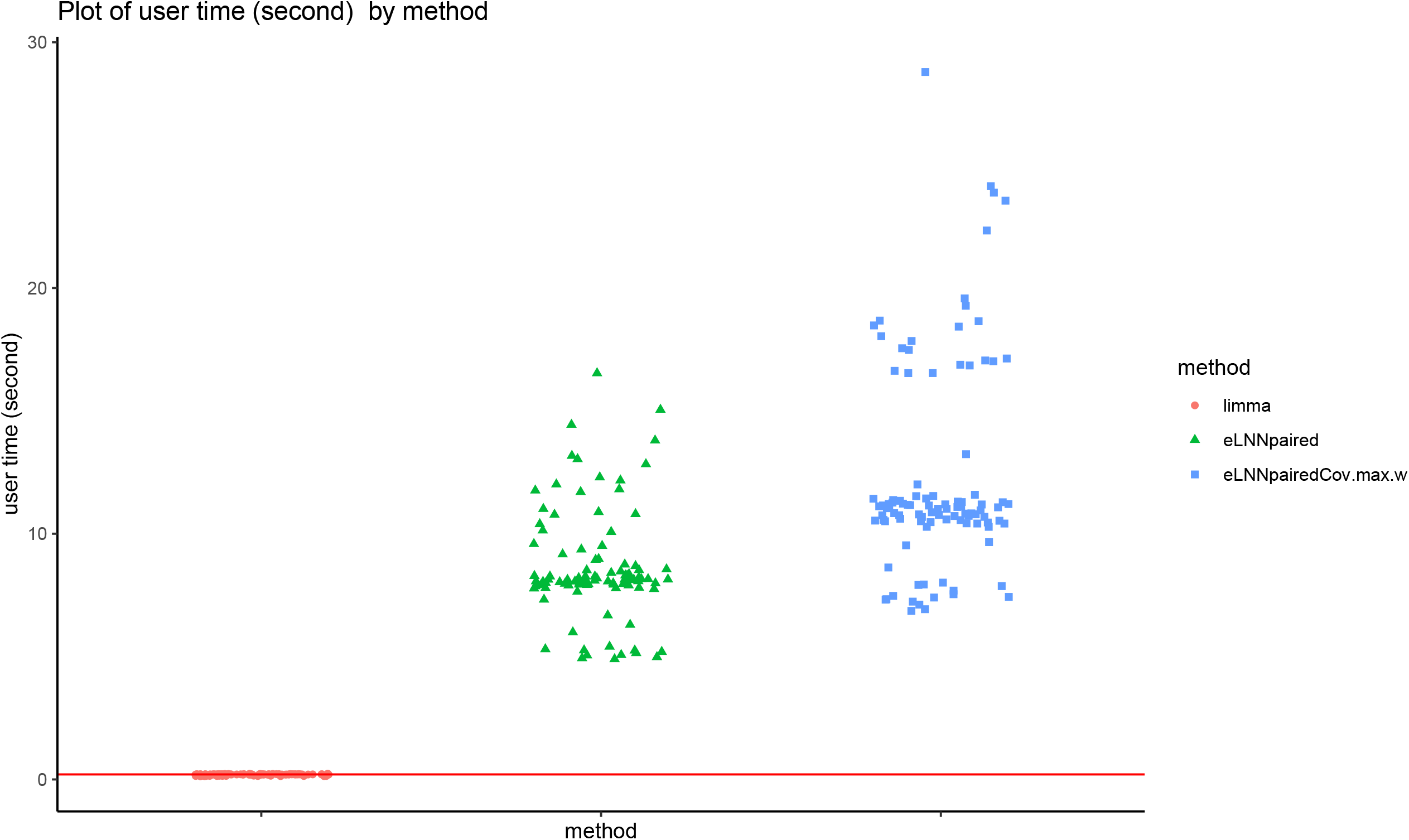

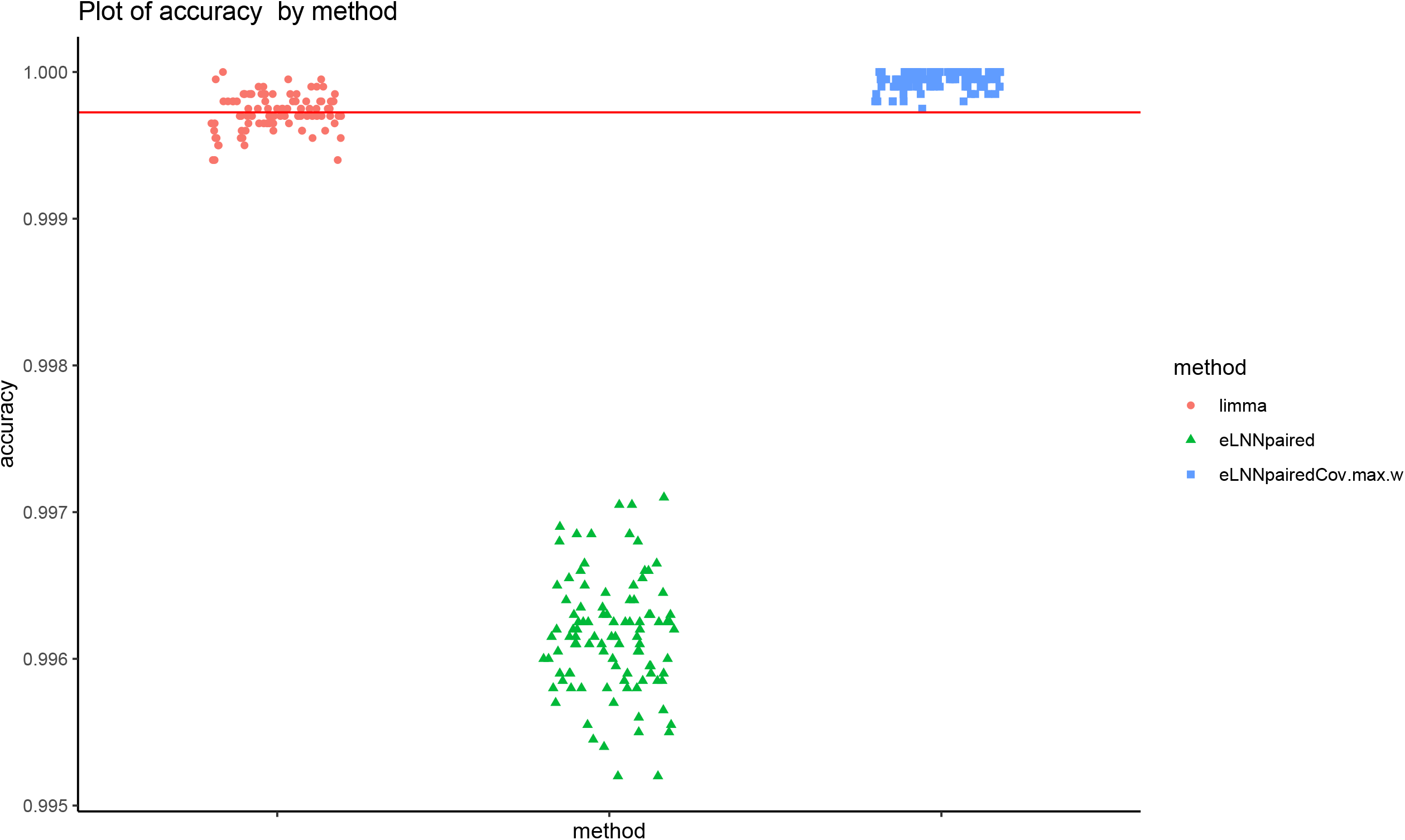

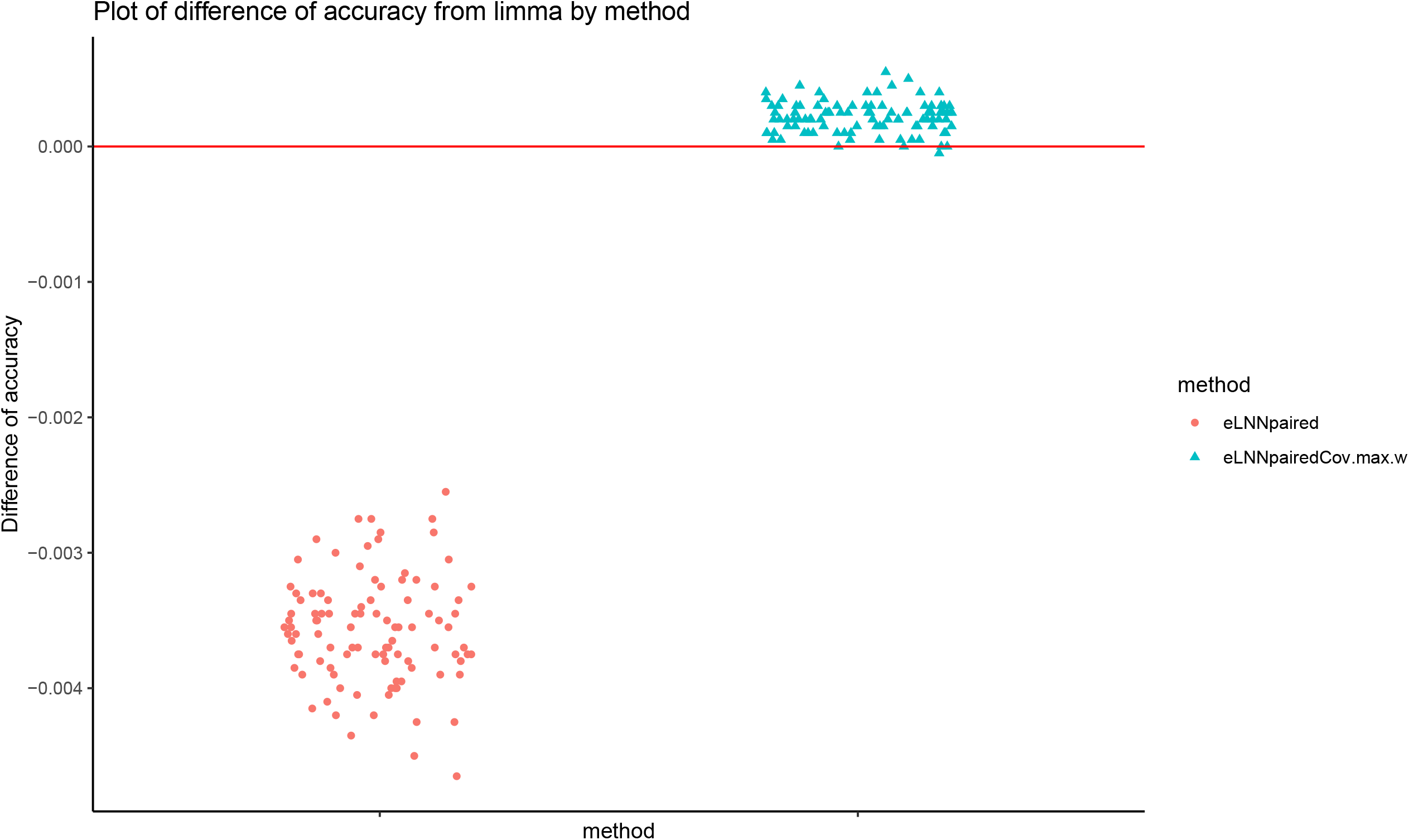

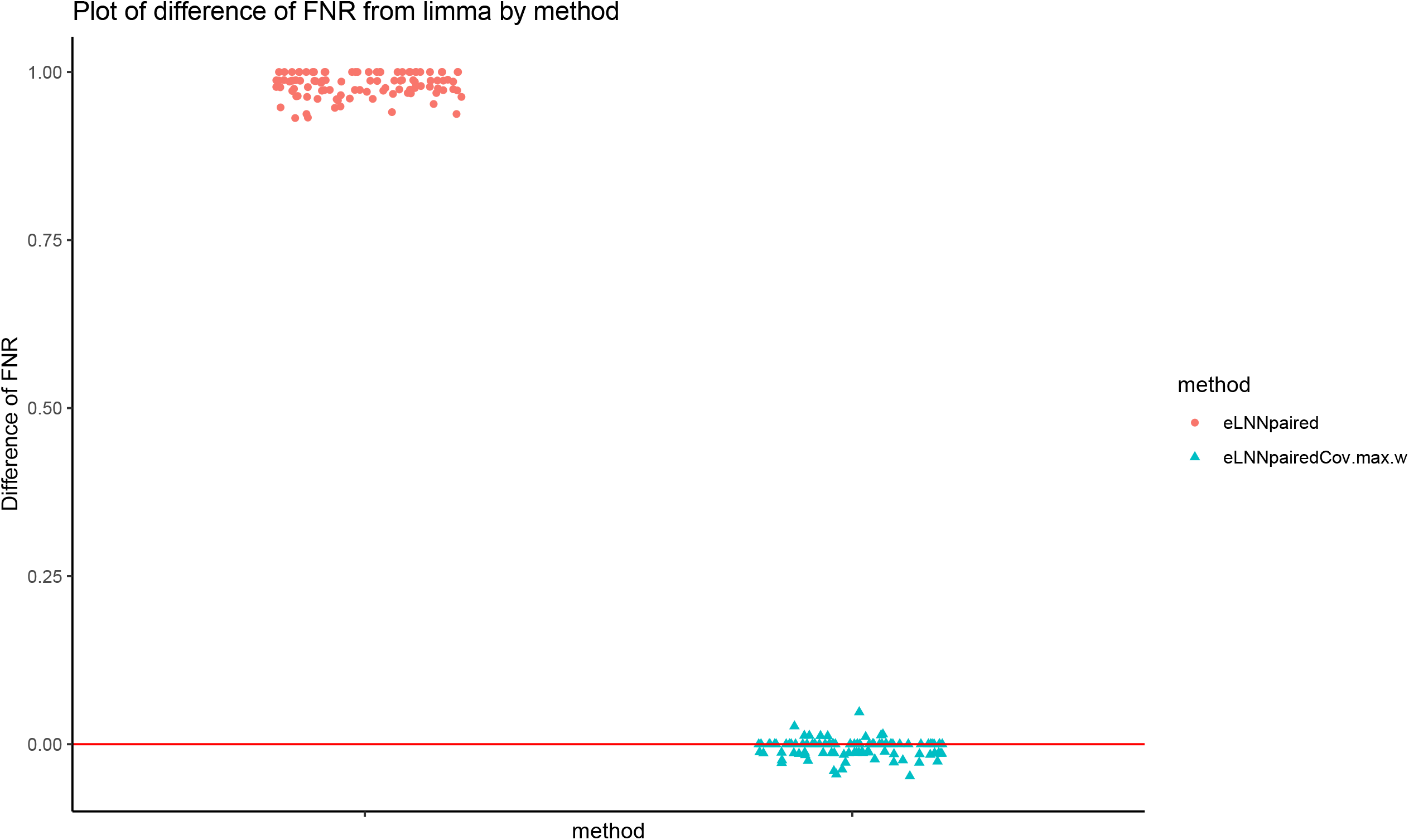

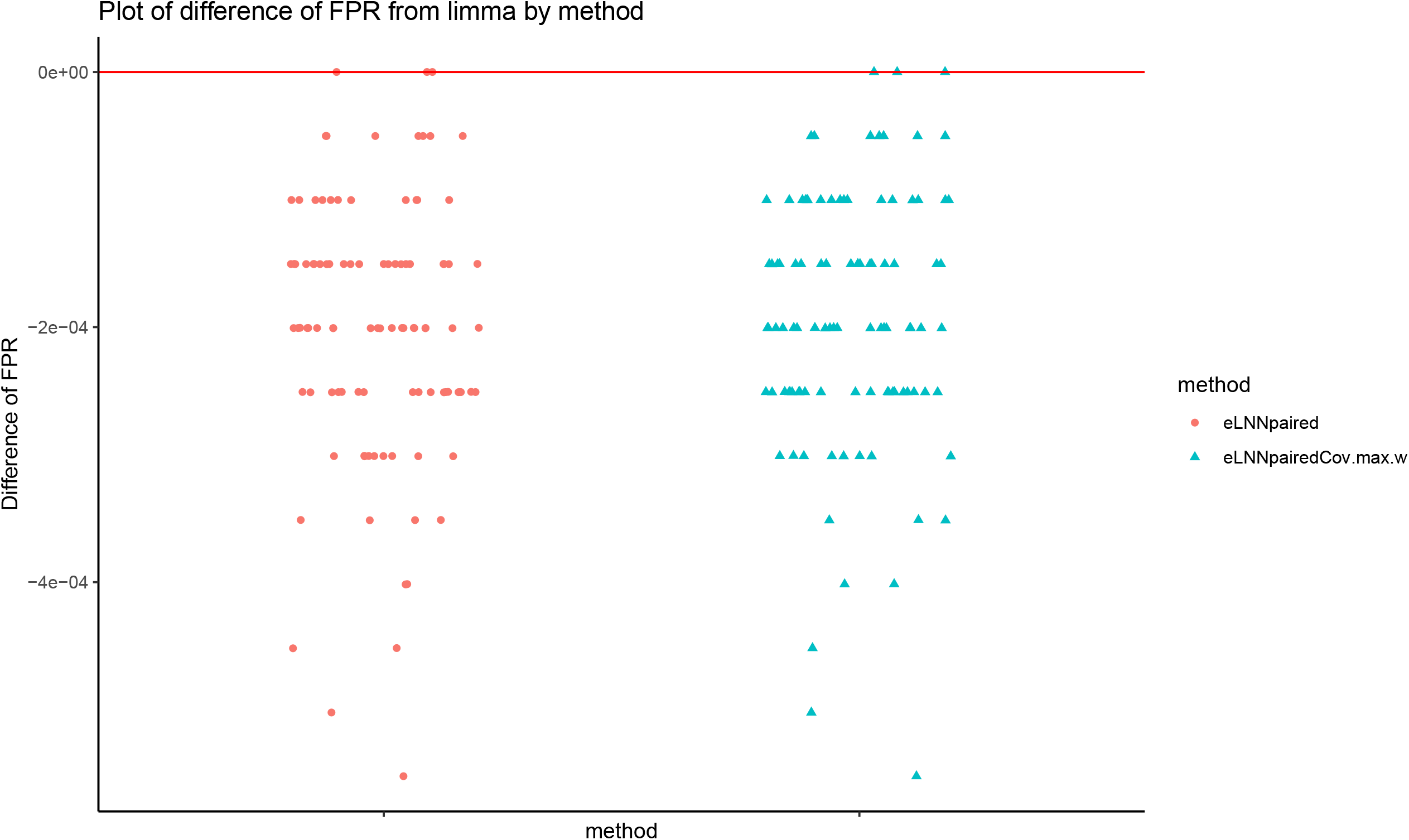

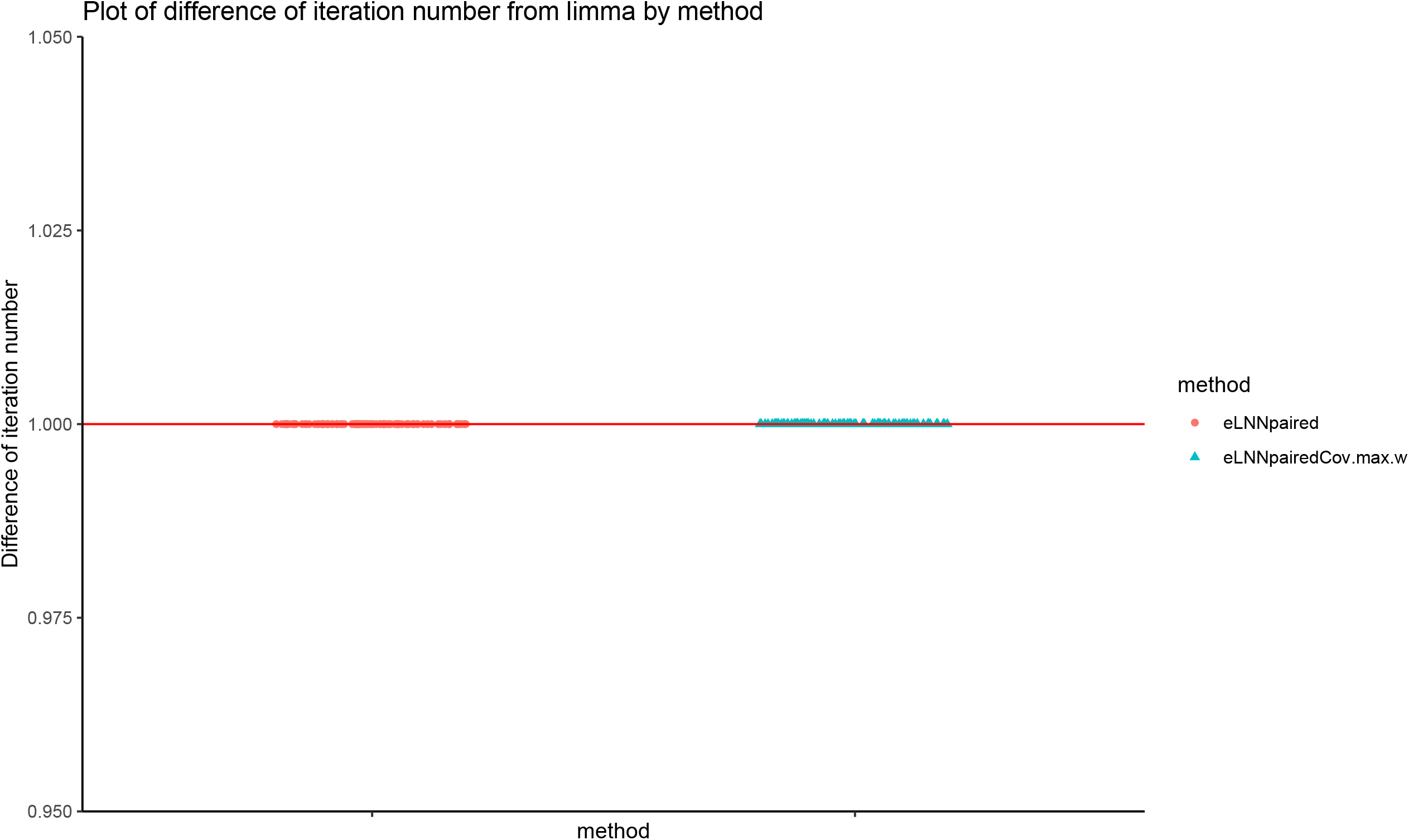

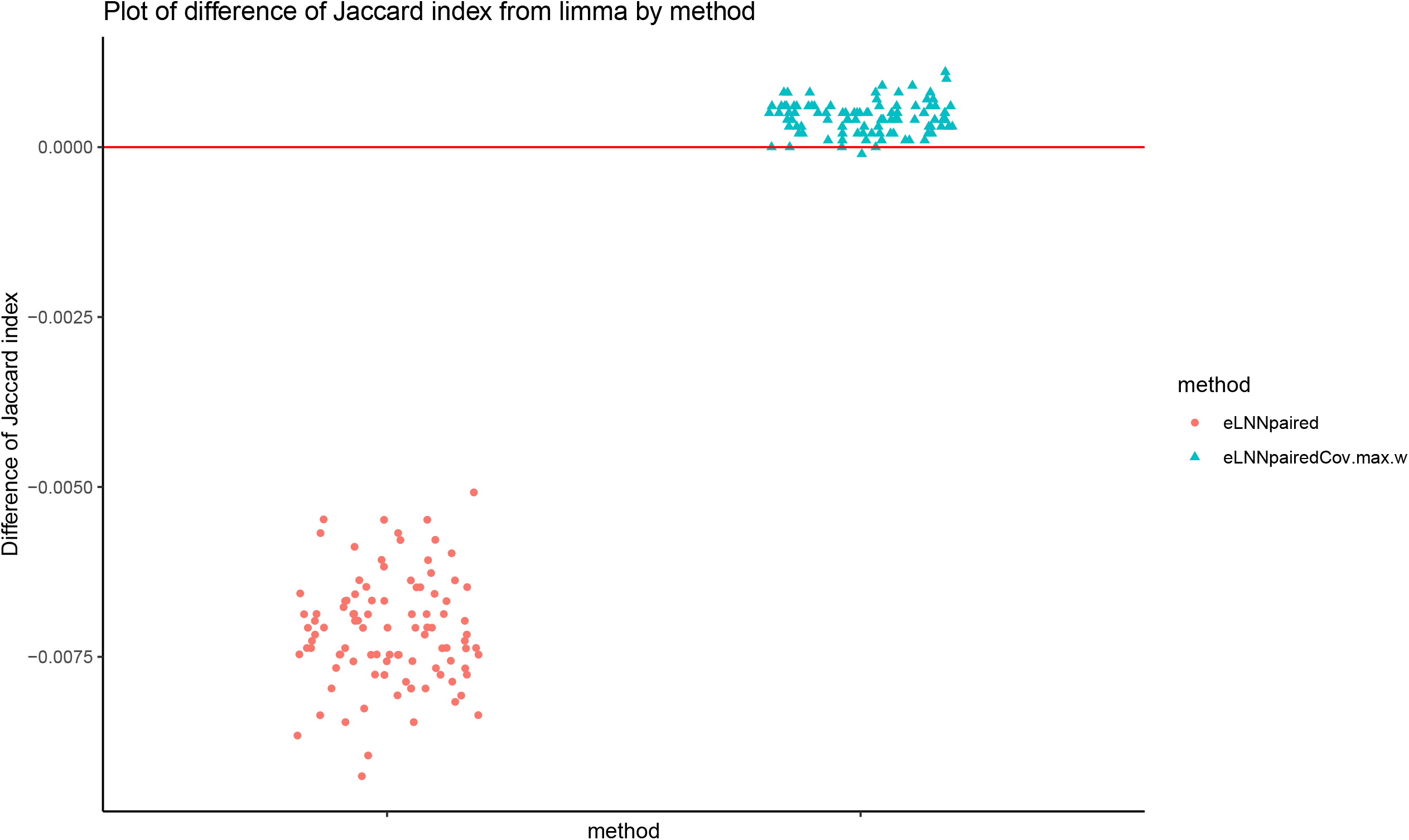

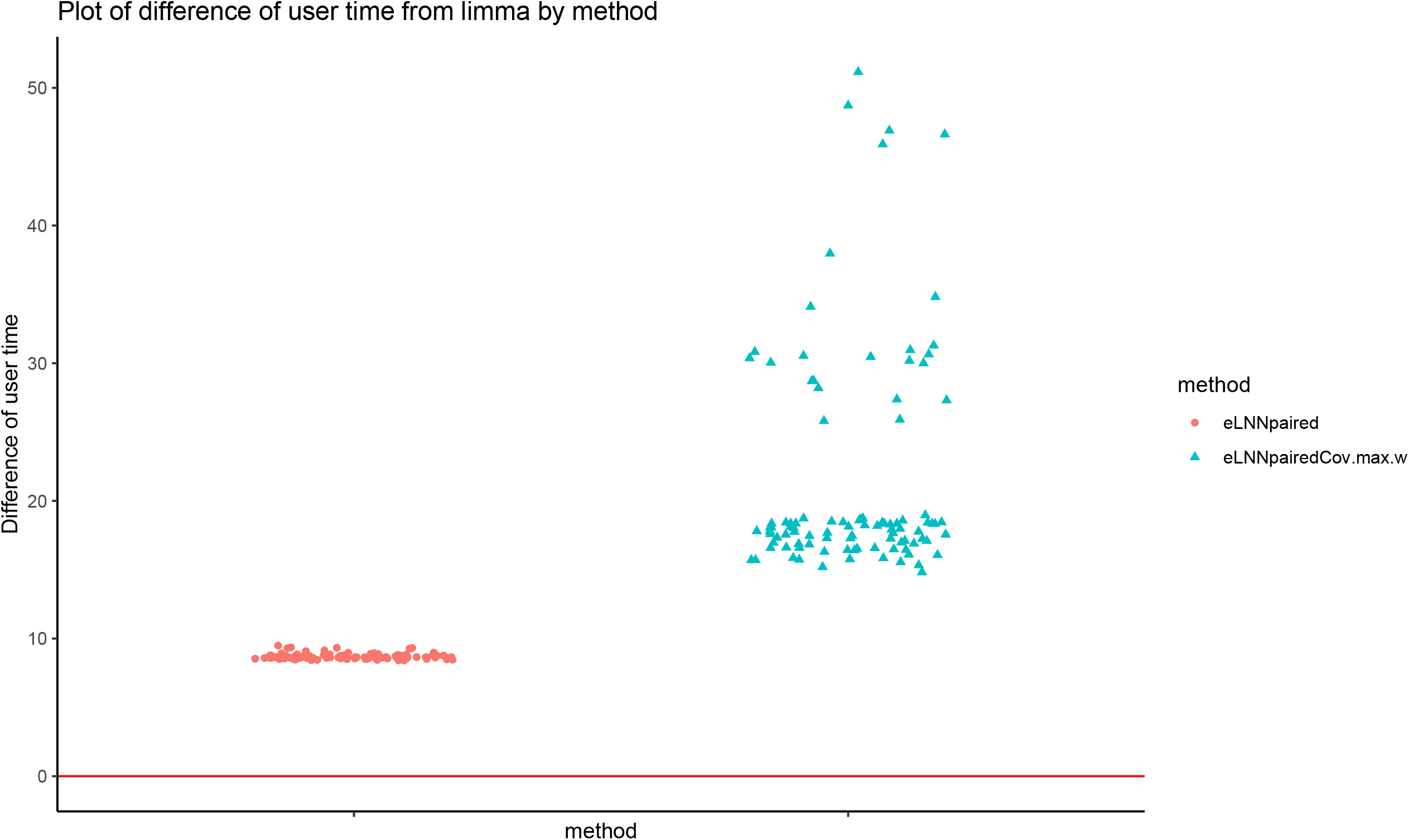

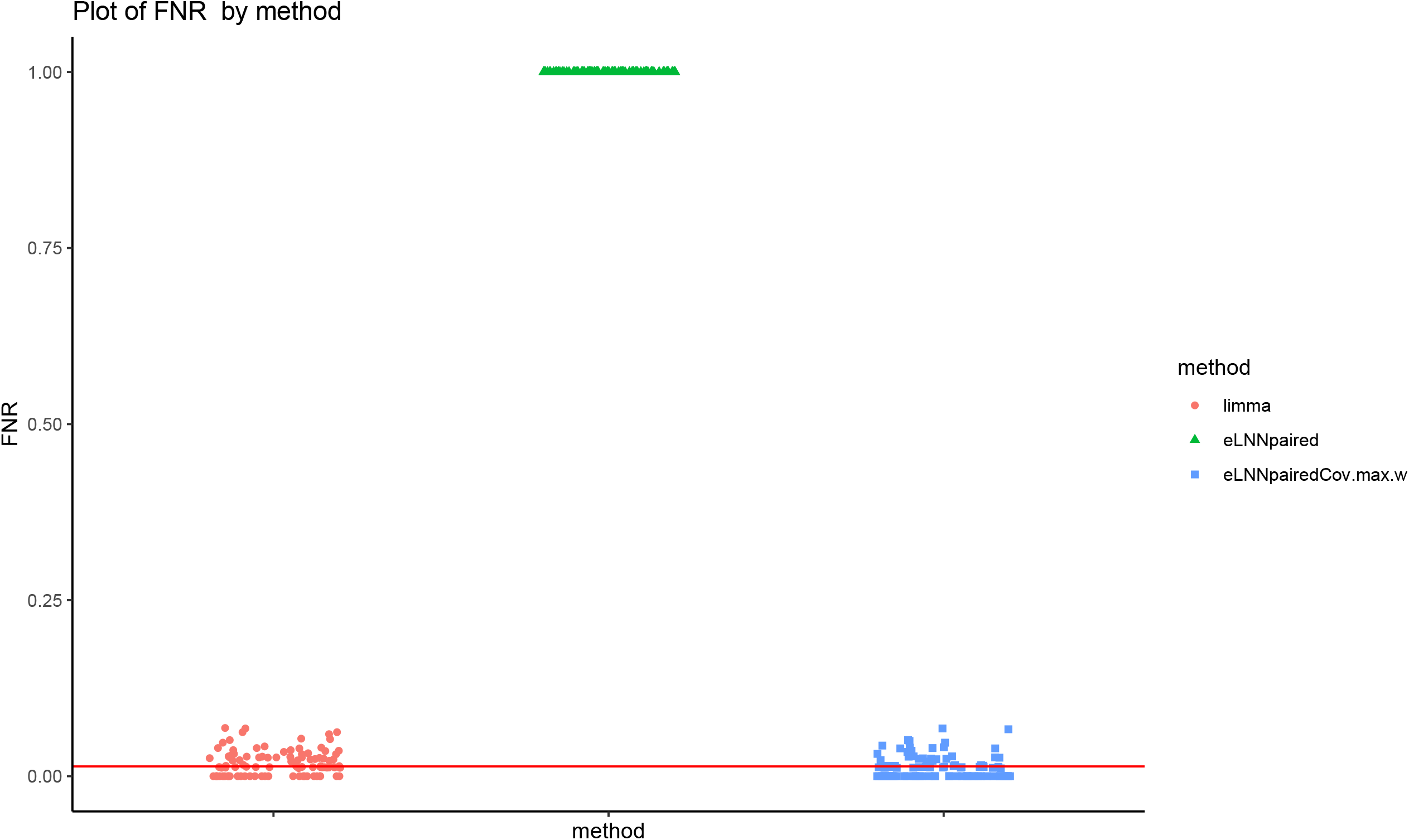

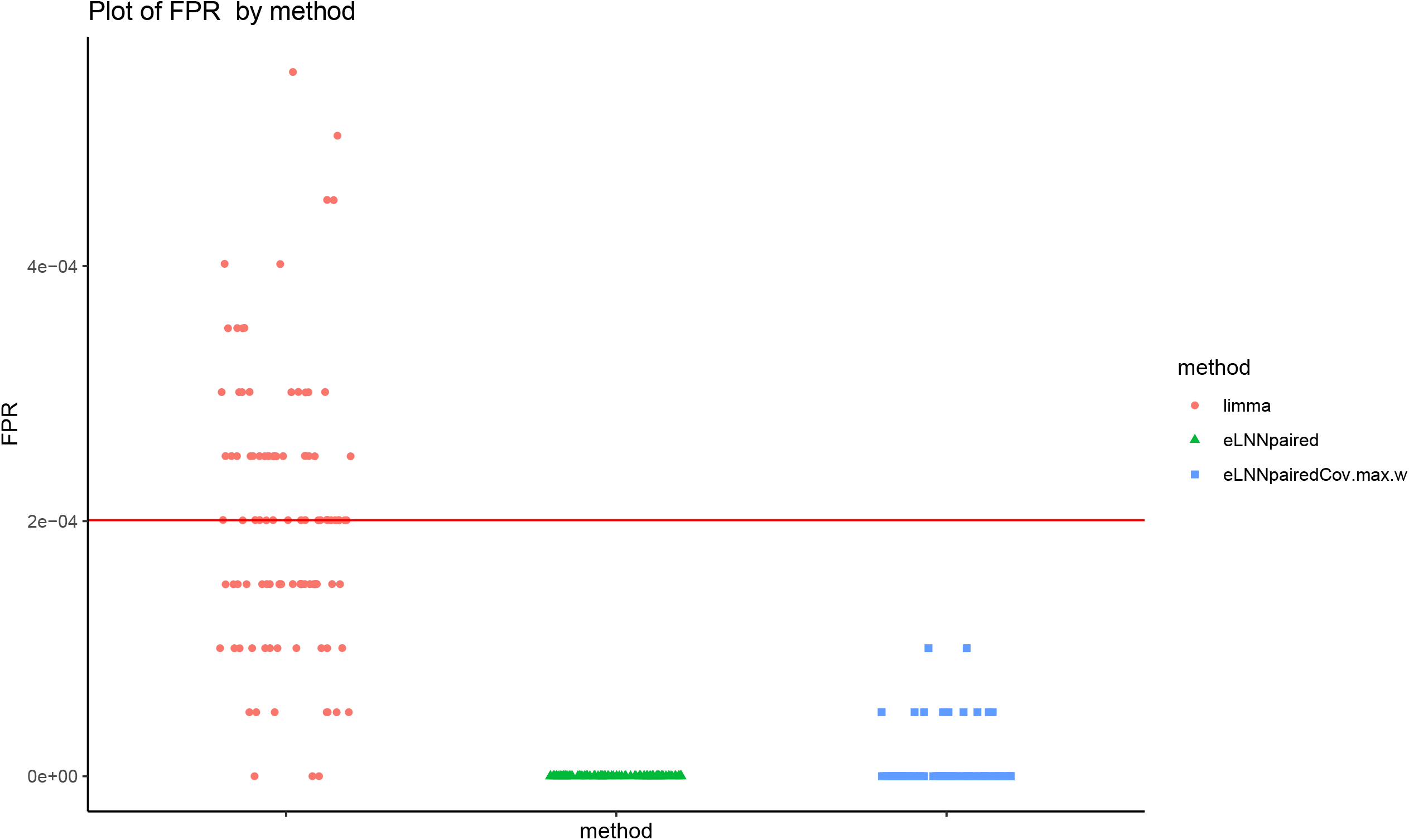

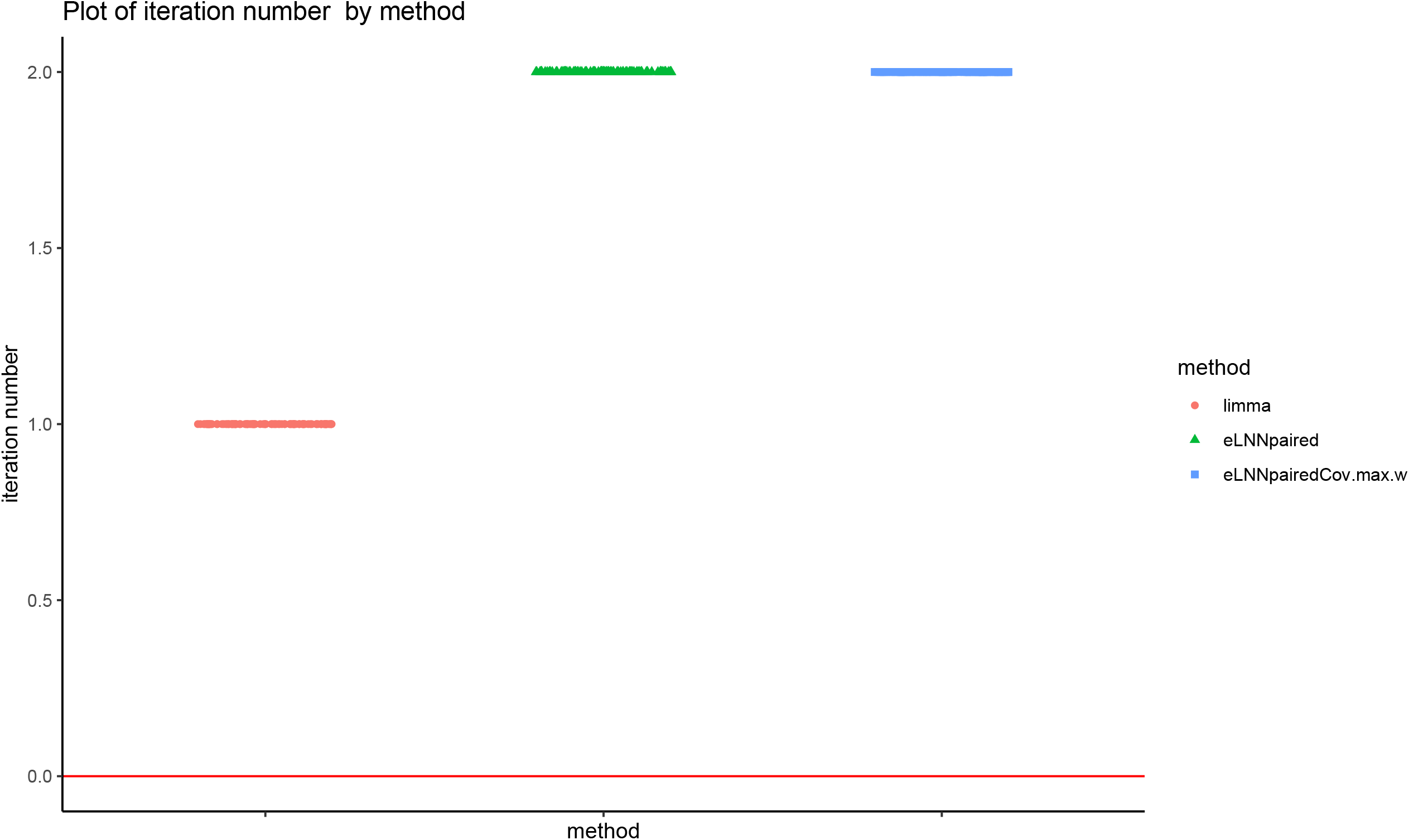

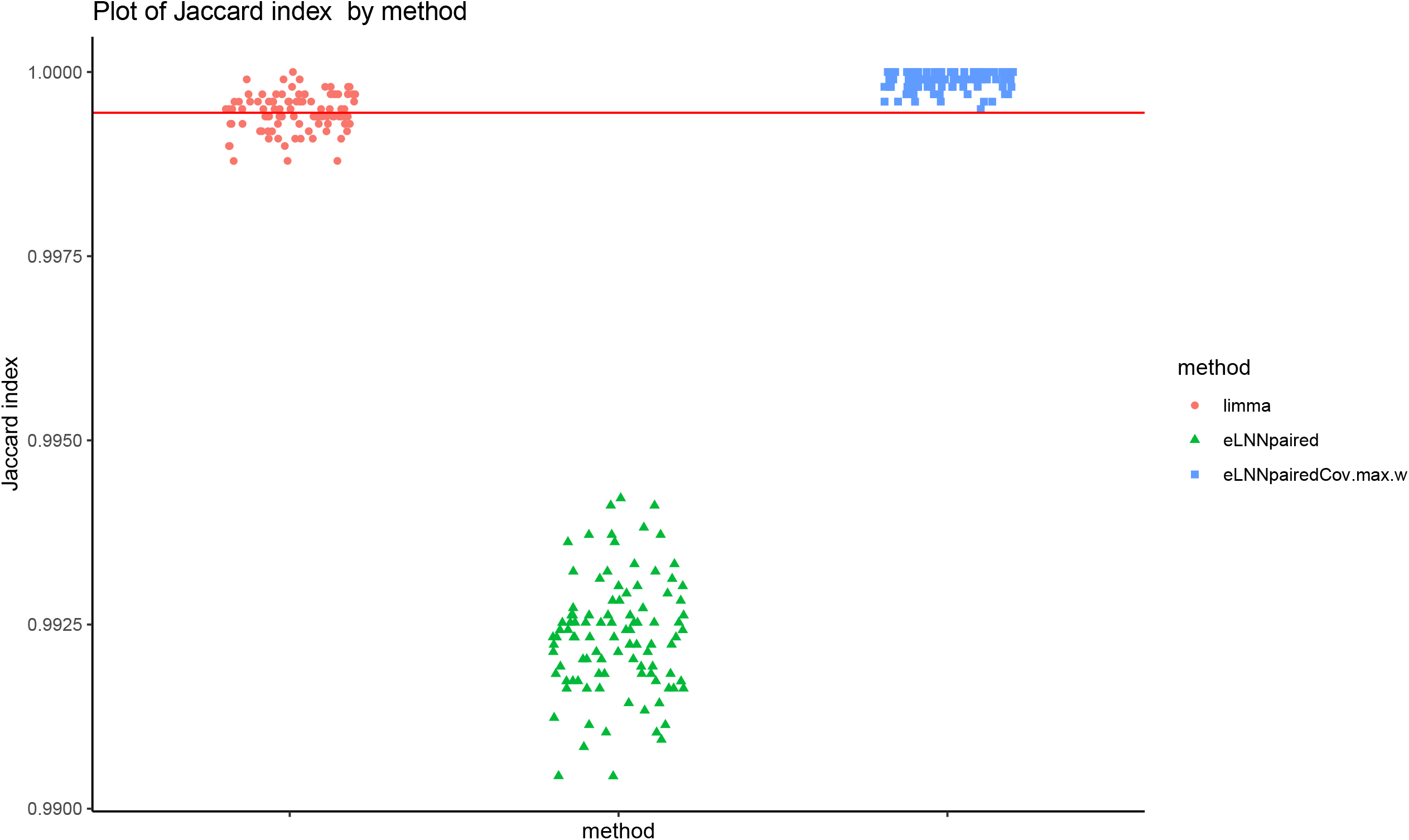

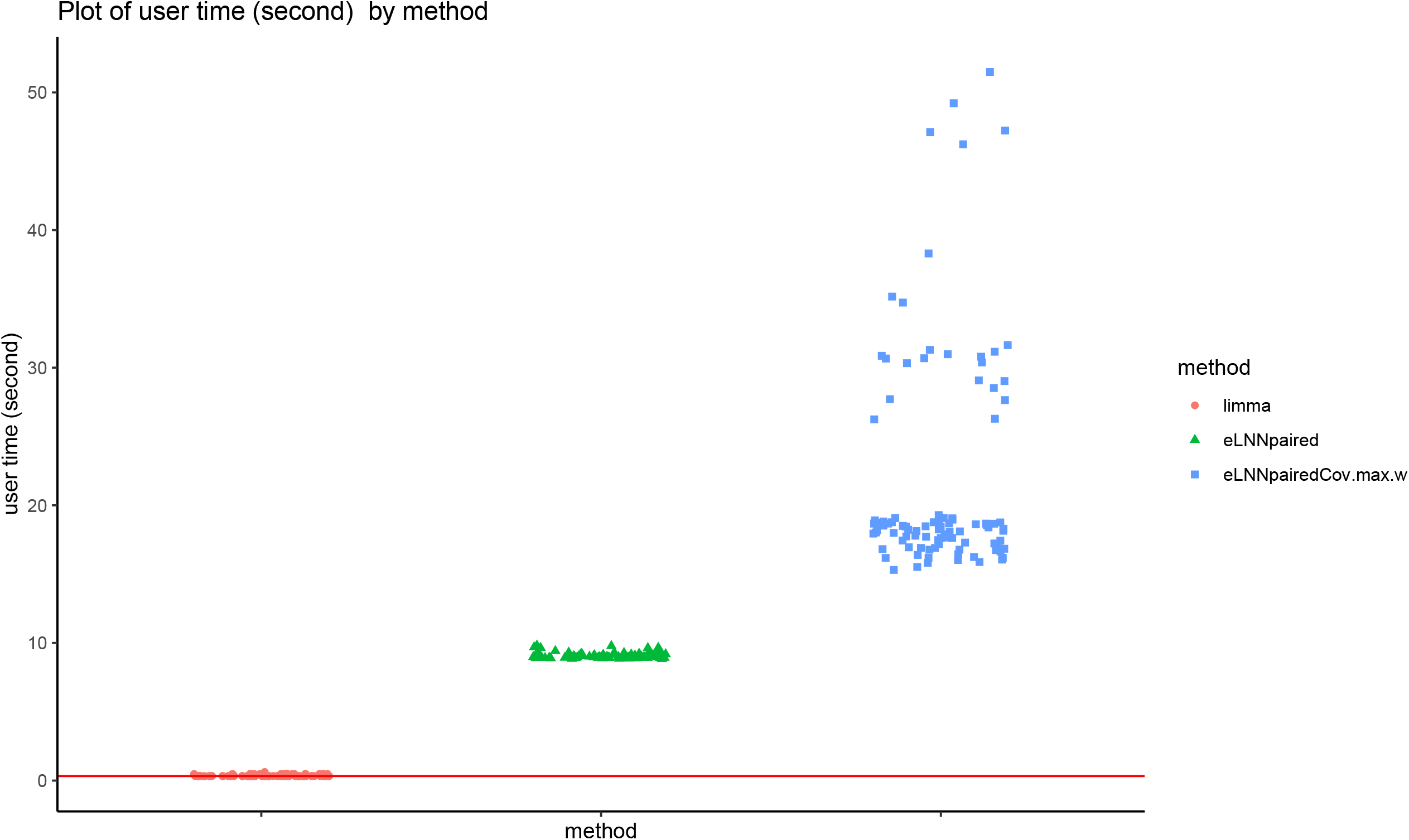

